# Immunoinformatic based analytics on T-cell epitope from spike protein of SARS-CoV-2 concerning Indian population

**DOI:** 10.1101/2021.01.07.425724

**Authors:** Sreevidya S. Devi, Manish Dwivedi

**Author notes:** Corresponding author, Email ID.

## Abstract

The whole world is drastically affected by the current pandemic due to severe virus, SARS-CoV-2 and scientists are rigorously looking for the efficient vaccine against it that become an emergent issue. Reverse vaccinology approach may provide us with significant therapeutic leads in this direction and further determination of T-cell / B-cell response to antigen. In the present study, we conducted population coverage analysis referring to the diverse Indian population. By using tools from Immune epitope database (IEDB), HLA- distribution analysis was performed to find the most promiscuous T-cell epitope out of *In silico* determined epitope of Spike protein from SARS-CoV-2. Selection of these epitopes have been conducted based on their binding affinity with the maximum number of HLA alleles belong to the highest population coverage rate values for the chosen geographical area in India. 404 cleavage sites within the 1288 amino acids sequence of spike glycoprotein were determined by NetChop proteasomal cleavage prediction suggesting that this protein has adequate sites in the protein sequence for cleaving into appropriate epitopes. For population coverage analysis, 221 selected epitopes are considered that shows the projected population coverage as 83.08% with 19.29 average hit (average number of epitope hits/HLA combinations recognized by the population) and 5.91 pc90 (minimum number of epitope hits/HLA combinations recognized by 90% of the population). 54 epitopes are found with the highest coverage among the Indian population and highly conserved within the given spike RBD domain sequence. Docking analysis of each epitope with their respective allele suggests that the epitope NSFTRGVYY represents highest binding affinity with docking score -7.6 kcal/mol with its allele HLA-C*07:01 among all the epitopes. Since the Covid-19 cases are still in progress and seem to remain like this until we find an effective vaccine, moreover in countries like India, vast diversity in the population may present a hindrance to particular vaccine efficiency. Outcomes from this study could be critical to design vaccine against SARS-CoV-2 for a different set of the population within the country.

## 1. Introduction

The immune system is an organization of cells and molecules with specialized roles in defending against infection [1]. The immune system functions as the body‘s resistance against virulent organisms and further unfamiliar agents [2]. T cell epitopes are short linear peptides that are either cleaved from antigenic proteins or generated by protein splicing [3]. T cell epitopes are presented in the surface of Major Histocompatibility Complex (MHC) proteins and in case of humans on class I or class II Human Leukocyte Antigen (HLA) molecules [2]. Epitope presentation depends on both MHC-peptide binding and T cell receptor (TCR) interactions [4,5]. MHC proteins are highly polymorphic and bind to a restricted set of peptides [2]. Consequently, a specific combination of MHC alleles present in a host restricts the choice of potential epitopes identified throughout an infection [2]. The conformation of a T cell epitope implanted in an MHC protein is significant for TCR recognition [6,7]. B cell epitopes on the other hand can be linear contiguous or discontinuous amino acid sequences that are brought together spatially in folded proteins [8].

As immune system is well-thought-out as a system of thousands of molecules, which directs to several intertwined responses, it is structurally and functionally distinct and this diversity differs both between individuals and temporally within individuals as a result it can create huge amounts of data [9]. The major advantage of immunoinformatics is that it can reduce the time and cost essential for laboratory analysis of pathogenic gene products. So, this information enables an immunologist to explore the potential binding sites, which, in turn, guides to the development of novel vaccines. This methodology is labelled as reverse vaccinology. Vaccination is an extremely successful approach to disease control in human and veterinary health care [10].

It examines the pathogen genome in order to identify potential antigenic proteins. This is beneficial as conventional approaches need to cultivate pathogen and then extract its antigenic proteins for analysis [11]. Immunoinformatics can also recognize virulence genes and surface□associated proteins.

The 2019 novel coronavirus (2019-nCoV) infection was emerged in Wuhan, China, in December 2019 and has rapidly multiplied throughout China and several other countries [12]. It was found that this SARS-CoV-2 virus has genome similarity with SARS□CoV up to 79.5% which has already caused a global epidemic in around 25 countries during 2002-2003 with 8096 confirmed cases and 96% with bat coronavirus [13, 14]. The virus has a positive□sense single□stranded RNA as their genetic constituent. It belongs to the family Coronaviridae of order Nidovirales and is a β□coronavirus [12]. The epidemiology of coronavirus demonstrates that human□to□human transmission of the virus happens through various routes like sneezes, cough, and respiratory droplets [12]. After days of illness, spread of SARS-CoV occurred widely which is associated with modest viral loads in the respiratory tract during initial stage of the illness, in later stages viral loads increases in approximately 10 days after the onset of symptoms. Epidemiologic examination in Wuhan city China recognized an early correlation with a seafood market where most affected people had worked or visited [15, 16, 17]. Numerous confirmed cases were registered rapidly all over the world as the outbreak progressed. As a result, the disease is marked as pandemic and was named as COVID□19. The disease was marked with extensive morbidity and mortality all over the world.

As the disease was spreading all over India initially in an uncontrollable manner, the necessity for developing an effective peptide vaccine component against the SARS□COV□2 was rising [12]. Our work was to discover appropriate T-cell epitopes from SARS COV2 spike glycoprotein RBD (Receptor Binding Domain), that can produce sufficient immune response against the SARS□COV□2 infection. By using immunoinformatics techniques, we could identify and characterize potential T□cell epitopes for the development of the epitopic vaccine against SARS□COV□2. The spike glycoprotein of SARS□COV□2 is chosen as the target as it produces a distinctive crown around the virus that projects from the viral envelope that helps in viral attachment with the host receptor [18]. Several immunoinformatic□based servers and software were used to explore spike glycoprotein to recognize many epitopes for an effective vaccine [19]. In order to identify the T cell epitopes that are effective to produce immune response among Indian population, we carried out a population coverage analysis within the Indian Population from the given MHC alleles and obtained a set of epitopes that is estimated to provide broad coverage within the population. Our findings provide a screened set of epitopes that may help to guide in experimental efforts towards the development of vaccines against SARS-CoV-2 for Indian population.

## 2. Materials and Methods

### 2.1 Sequence Retrieval

Protein sequence of prefusion SARS-COV-2 spike glycoprotein with an RDB (NCBI ID-6VSB_A) was retrieved from NCBI database (https://www.ncbi.nlm.nih.gov/protein/6vsb_A) [20]. NCBI‘s Protein resources comprise protein sequences and structures and associated comparison and visualization tools, as well as databases and tools to predict and analyse functional domains. Among the different spike glycoprotein sequences available in the database, chain A of spike glycoprotein with RBD domain is considered as it plays a huge role in interacting with the human ACE2 receptor. The spike glycoprotein sequence in FASTA format is downloaded.

## 2.2. Antigenicity Prediction

Antigenicity prediction of the protein sequence is significant as it gives an idea about the possibility of the viral protein sequence to be recognized by the immunogenic cells present in the human body.

Vaxijen v2.0 server [21] (http://www.ddg-pharmfac.net/vaxijen/VaxiJen/VaxiJen.html) is used to predict antigenicity as it is the initial server for alignment-independent prediction of protective antigens This is alignment independent predictor based on auto-cross covariance (ACC) transformation epitopes sequences into uniform vectors of principle amino acid properties. Accuracy of this server varies in between 70 to 89% depending on targeted organism. This software requires FASTA submitted amino acid sequences and was established to certificate antigen classification uniquely based on the physicochemical properties of proteins without recourse to sequence alignment. The default parameters (threshold=0.4, ACC output) were used in contrary to viral species to predict the antigenicity of full-length protein sequence.

### 2.3. Physio-Chemical Parameter Prediction of Sequence

Protparam online tool (https://web.expasy.org/protparam/) [22] from Expasy is used to predict the physio-chemical parameter of the viral protein sequence. This tool assists to compute physio-chemical characters of primary protein structure such as molecular weight, theoretical pI, amino acid composition, atomic composition, extinction coefficient [23], estimated half-life [24], instability index [25], aliphatic index [26], and grand average of hydropathicity (GRAVY) [27]. A stable protein sequence with low molecular weight is considered to be decent for epitope prediction.

### 2.4. Super Secondary Protein Prediction

Super secondary protein prediction is used to classify the total number of domains present in the protein sequence as a greater number of domains is truly decent for an improved antigen and to design an epitope-based vaccine. NCBI Conserved Domain Database (CDD) [28] (https://www.ncbi.nlm.nih.gov/Structure/cdd/wrpsb.cgi) is used to predict the domain present in the protein sequence as The Conserved Domain Database is a database of well-annotated multiple sequence alignment models and derived database search models, for ancient domains and full-length proteins. Functionally important sites were defined as those residues making an interaction with a ligand or a macromolecule. CDD alignments signify alignments of conserved core structures formed by presumably homologous sites, and positions exterior to the conserved cores are removed from the alignment. Fasta format of protein sequence is specified as input and searching is done against CDD v3.18 which is a superset including NCBI-curated domains and data imported from Pfam [29], SMART [30], COG [31], PRK, and TIGRFAM [32]. All other parameters are set into default for calculation.

### 2.5. Proteasomal Cleavage Prediction

The generation of cytotoxic T lymphocyte (CTL) epitopes from an antigenic sequence comprises of number of intracellular processes, including production of peptide fragments by proteasome (proteasomal cleavage) and transport of peptides to endoplasmic reticulum through transporter associated with antigen processing (TAP). The NetChop 3.1 server (http://www.cbs.dtu.dk/services/NetChop/) [33] is used to predict the proteasomal cleavage of peptide sequence in the host which produces neural network predictions for cleavage sites of the human proteasome. Protein sequence in FASTA format is given as input by keeping the threshold into 0.5 and method as C term 3.0.

### 2.6. Identification of HLA Alleles

The Allele Frequency Net Database (AFNDB) (http://www.allelefrequencies.net/hla6006a.asp) [34] is used to identify the HLA allele frequency of the population. This database grants the scientific community with a freely accessible repository for the storage of frequency data (alleles, genes, haplotypes and genotypes) related to human leukocyte antigens (HLA), killer-cell immunoglobulin-like receptors (KIR), major histocompatibility complex Class I chain related genes (MIC) and a numerous cytokine gene polymorphism in worldwide populations. At present, AFND contains >1600 populations from >10 million healthy individuals, making AFND aa appreciated source for the analysis of some of the most polymorphic regions in the human genome. Identification of HLA classical allele frequency of all loci from all geographical regions was carried out by selecting country as India and results are shown by sorting allele frequency from highest to lowest. Alleles are obtained from all sources covering Literatures, Proceedings of IHWs and Unpublished.

### 2.7. MHC Class-I T-Cell Epitope Prediction

The primary step on applying bioinformatics to epitope based vaccine development comprises of differentiating epitopes that are potentially immunoprotected from epitopes that are not. Since T-cell epitopes are connected in a linear form to MHCs, the interface between ligands and T-cells can be modelled with accuracy. MHC-I binding predictors are presently very resourceful and have extensive allelic coverage, a prediction accuracy in the range of 90–95% positive prediction value has been estimated. Several T-cell epitope prediction tools based on diverse algorithms are employed as it aids to recognize the finest 9-mer epitopes from the protein sequence. Most of the prediction server‘s take protein sequence in plain text format expect few. HLA alleles are selected on the basis of their allele frequency and percentage of individuals that have the allele. HLA alleles having percentage above 9 are selected. Amongst the several servers for MHC-I alleles, RANKPEP (http://imed.med.ucm.es/Tools/rankpep.html) [35], which foretells peptide binders to MHC-I and MHC-II molecules from protein sequences or sequence alignments using Position Specific Scoring Matrices (PSSMs). Moreover, it predicts those MHC-I ligands whose C-terminal end is likely to be the outcome of proteasomal cleavage. Protein sequence in FASTA text format is given as input by keeping all the parameters as default. It was used to predict epitopes binding to HLA-A*01:01, HLA-A*02:01, HLA – A*02:06, HLA-A*03:01, HLA-B*07:02, HLA-B*35:01,HLA-B*51:01,HLA-A*29:02,HLA-A*11:01,HLA-A*68:01,HLA-B*58:01 and HLA-B*57:01. SYFPEITHI(http://www.syfpeithi.de/bin/MHCServer.dll/EpitopePrediction.htm) [36] is another class I T-cell epitope prediction tool that utilities published motifs to predict the appropriate epitopes. Epitopes binding to HLA-A*11:01, HLA-A*68:01, HLA-B*07:02 and HLA-B*50:01 are predicted by means of this tool. A neural-network based MHC Class I binding peptide prediction tool called ANNPRED (https://webs.iiitd.edu.in/raghava/nhlapred/neural.html) [37] which utilizes artificial neural networks to predict HLA-A*02:06, HLA-A*03:01 and HLA-A*11:01 binding epitopes are also used. Additional neural-network based tool called COMPRED (https://webs.iiitd.edu.in/raghava/nhlapred/comp.html) [38] which is a hybrid method of artificial neural networks and quantitative matrices are used to predict epitopes binding to HLA-A*02:06, HLA-A*03:01 and HLA-A*11:01 alleles. Both the server‘s predicted the epitope from default parameters. T cell epitope processing prediction tools of IEDB Analysis Resource (http://tools.iedb.org/mhci/) [39] that will take in an amino acid sequence, or number of sequences and govern each subsequence‘s ability to bind to a definite MHC class I molecule, is used to predict epitopes interacting with HLA-A*01:01, HLA-A*02:06, HLA-A*03:01, HLA-A*011:01, HLA-A*29:02, HLA-A*68:01, HLA-B*07:02, HLA-B*18:01, HLA-B*35:01, HLA-B*35:03,HLA-B*51:01,HLA-B*57:01,HLA-B*58:01,HLA-C*04:01,HLA-C*07:01,HLA-C-07:02 alleles. The prediction method was IEDB recommended which is the default prediction method selection. MHC source species are selected as humans and the output is sorted based on predicted IC50 values. Epijen v1.0 (http://www.ddg-pharmfac.net/epijen/EpiJen/EpiJen.htm) [40], is a reliable multi-step algorithm for T cell epitope prediction, which belongs to the next generation of in silico T cell epitope identification methods. These methods aim to reduce subsequent experimental work by improving the success rate of epitope prediction. This is used to predict class I T-cell epitopes binding to HLA-A*01:01, HLA-A*02:06, HLA*A-03:01, HLA*A-11:01, HLA-A*68:01, HLA-B*35:01 by keeping all parameters as default. NetMHCpan 4.1 server (http://www.cbs.dtu.dk/services/NetMHCpan/) [41] is also an artificial neural network based MHC prediction tool skilled on a combination of more than 850,000 quantifiable Binding Affinity (BA) and Mass-Spectrometry Eluted Ligands (EL) peptides to predict the binding of peptides to any MHC molecule of known sequence. Class I epitopes interacting with HLA-A*01:01, HLA-A*03:01, HLA-B*07:02 and HLA-B*58:01 are predicted. Input type is FASTA format and all other parameters are kept default for the prediction of 9-mer epitopes.

### 2.8. Population Coverage

Population Coverage tool of IEDB (http://tools.iedb.org/population/) [42] is used as it calculates the fraction of individuals predicted to respond to a given set of epitopes with known MHC restrictions. This calculation is made based on HLA genotypic frequencies assuming non-linkage disequilibrium between HLA loci. The most prominent computationally validated epitopes predicted from several epitope prediction tools were taken into consideration to predict the respond of individuals form Indian population towards the predicted epitopes. 221 philandering high or moderate immunogenic T-cell epitopes having the predicted reputed epitopic core sequences identified from various epitope prediction tools along with its corresponding class I HLA alleles are submitted to the tool to identify the epitopic coverage of Indian population by keeping other default parameters on.

### 2.9. Epitope Conservancy and Immunogenicity Prediction

In an epitope-based vaccine design, the use of conserved epitopes would likely provide wider protection across multiple strains, or even species, as compared to epitopes obtained from extremely variable genome regions. The conservancy tool was designed to analyse the variability or conservation of epitopes. This tool calculates the degree of conservancy of a set of peptide/epitope sequences within a given set of protein sequences. Epitope Conservancy Analysis of IEDB Analysis Resource (http://tools.iedb.org/conservancy/) [43] computes the degree of conservancy of an epitope within a given protein sequence set at a given identity level. 54 selected epitopes showing high coverage from population coverage analysis tool is selected and are given as plain text as input. Along with that protein sequence in FASTA format is also given by keeping other parameters as default. To predict the degree of immunogenicity of the selected peptides, another tool from IEDB Analysis Resource named as Class I Immunogenicity (http://tools.iedb.org/immunogenicity/) [44] is employed. This tool utilizes amino acid properties as well as their position within the peptide to predict the immunogenicity of a peptide MHC (pMHC) complex. The selected 54 epitopes are given as input and other parameters are given as default.

### 2.10. Tap Binding Predictions

TAP refers to transport of peptides to endoplasmic reticulum through transporter associated with antigen processing. TAP prediction is essential to identify whether the predicted epitopes will be successfully transported to endoplasmic reticulum as it is significant for the epitope to produce a response in the human body. All those epitopes with vital IC50 values can be chosen for epitope-based vaccine design. The understanding of selectivity and specificity of TAP may contribute significantly to prediction of the *MHC* class I restricted T-cell epitopes. For predicting binding affinities of peptides to TAP, an online tool TAPREG (http://imed.med.ucm.es/Tools/tapreg/) [45] from Immunomedicine Group is used. It predicts the binding affinity by using a Support Vector Machine model. Selected 54 epitopes in FASTA format is given as input by keeping all other parameters as default.

### 2.11. Allergy Prediction

AllerTOP v. 2.0 (https://www.ddg-pharmfac.net/AllerTOP/feedback.py) [46], a Bioinformatics tool for allergenicity prediction is used to anticipate whether the determined epitopes from viral protein sequence would produce an allergic response on human body. AllerTOP is the primary alignment-free server for in silico prediction of allergens built on the key physicochemical properties of proteins. Selected 54 epitopes in plain text is given as input to compute the allergenicity of the sequence. Significantly, as well allergenicity AllerTOP also envisages the most probable route of exposure. In contrast to other servers for allergen prediction, AllerTOP outperforms them with 94% sensitivity.

### 2.12. Toxicity Extrapolation

The toxicity of selected epitopes are predicted using ToxinPred (https://webs.iiitd.edu.in/raghava/toxinpred/multi_submit.php) [47] web server. This tool permits the users to identify extremely toxic or non-toxic peptides from large number peptides offered by a user. It predicts their toxicity along with all the vital physio-chemical properties like hydrophobicity, charge pI etc. of peptides offered by users. This method was developed based on the machine learning technique and quantitative matrix using different properties of peptides. 54 selected peptides are given in FASTA format as input and all other parameters are kept default.

### 2.13. Epitope 3-D Structure Prediction and Ramachandran Plot Analysis

Structure prediction of epitopes can be useful to design epitope-based vaccines against spike protein of SARS-COV2. PEP-FOLD server (https://mobyle.rpbs.univ-paris-diderot.fr/cgi-bin/portal.py#forms::PEP-FOLD3) [48] which is based on a new de novo approach to predict 3D peptide structures from sequence information is used to predict the epitope structure. PEP-FOLD is based on the concept of structural alphabet (SA) and uses a Hidden Markov Model (HMM)-derived SA of 27 letters to describe proteins as series of overlapping fragments of four amino acids [49]. PEP-FOLD uses a two-step procedure: prediction of a limited set of SA letters at each position from sequence, and then assembly of the prototype fragments associated with each SA letter using a revised version of our greedy algorithm [50, 51] and a generic protein coarse-grained force field. 5 selected 9-mer epitopes showing highest population coverage as well as decent TAP prediction values are given to predict its 3-D structure. PROCHECK server (https://servicesn.mbi.ucla.edu/PROCHECK/) [52] is used to analyse the Ramachandran plot of the spike protein structure. Ramachandran plot shows the number of amino acid residues in the allowed region and predict the validity of the protein model that can be used to indicate the high quality of models for each epitope.

### 2.14. Molecular Docking Analysis of Predicted Epitopes

Autodock Vina [53] is used to study the molecular interactions between predicted epitopes with their respective alleles. AutoDock is free of charge practices that have been extensively cited in the literature as vital tools in structure-based drug design. It can be used for both protein-ligand docking as well as protein-peptide docking. The peptides and proteins are modified using MGL tools. 3D structure of 5 9-mer epitopes NSFTRGVYY, ASFSTFKCY, TRFQTLLAL, RFDNPVLPF and TRFASVYAW predicted from PEP-FOLD server is converted into PDBQT format as Autodock Vina accepts only molecules in this format for docking. These peptides are then docked with their allele‘s HLA-C*07:01, HLA-A*11:01, HLA-A*01:01, HLA-C*04:01 respectively. Crystal structure of HLA alleles HLA-A*01:01 and HLA-A*11:01 is retrieved from PDB and all the HETATM and additional chains were removed using Discovery Studio. 3-D structure of alleles HLA-C*07:02 and HLA-C*04:01 were modelled using Swiss-Model and were refined using Galaxy tool. The results from Autodock Vina were visualized using Pymol and the interactions were analyzed using Discovery Studio.

## 3. Results

### 3.1. Antigenicity prediction

Vaxijen v2.0 analysis of prefusion spike glycoprotein with RDB in 0.4 threshold exhibited an antigenicity of 0.4512 which indicates that the selected sequence is a probable antigen. So, it confirms that the protein sequence can be well considered for the epitope prediction.

### 3.2. Physio-Chemical Parameter Prediction of Sequence

Physio-chemical parameter prediction of full-length protein sequence using Protparam online tool helped to identify characteristics important for epitope prediction from a protein sequence. The molecular weight is computed as 142274.61 with theoretical pI **=** 6.14. The estimated half-life is 30 hours for mammalian reticulocytes, (in vitro). The instability index (II) is computed to be 31.58 which classifies the protein as stable. The Aliphatic index is computed to be 81.58 with Grand average of hydropathicity (GRAVY) = −0.163.

### 3.3. Super Secondary Protein Prediction

Super secondary protein prediction with NCBI Conserved Domain Database (CDD) specifies that there are three domains present in the protein sequence. One amongst it is Corona_S2 super family (Coronavirus S2 glycoprotein) of length 601 seen between sequence interval 662-1208 with Bit score 794.41 and E-value 0e+00. The next is SARS-CoV-2_Spike_S1_RBD (receptor-binding domain of the S1 subunit of severe acute respiratory syndrome coronavirus 2 Spike (S) protein) presents in the sequence interval 319-541 of length: 223, Bit Score: 492.30 and E-value 9.12e-166. The third domain to be identified is Spike-COV-like_S1_NTD (N-terminal domain of the S1 subunit of the Spike (S) protein from Severe acute respiratory syndrome coronavirus and related beta coronaviruses in the B lineage) of length 280 present in the sequence interval 13-304 with bit score 458.72 and E-value 4.53e-152. Among the three domains recognized, SARS-CoV-2_Spike_S1_RBD is significant as it plays a key role in binding of SARS-COV2 with ACE2 receptor of human host. Entire sequences involved in respective domain were also identified as it aids to distinguish whether the predicted epitopes exist in the major domains.

### 3.4. Proteasomal Cleavage Prediction

MHC class I binding predictions are very accurate for most of the identified MHC alleles. However, these estimation could be additionally enhanced by integrating proteasome cleavage. NetChop proteasomal cleavage prediction prophesied that the spike glycoprotein sequence has 404 cleavage sites within 1288 amino acids. It specifies that there are adequate sites in the protein sequence for cleaving into appropriate epitopes.

### 3.5. Identification of HLA Alleles

Identification of HLA alleles with Allele Frequency Net Database (AFND) helped to recognize the class I alleles commonly seen in the Indian population based on their Allele Frequency and Percentage of Individuals that have the allele. Alleles are identified from all the regions of Indian population. The results consist of Allele, Population from which alleles are identified along with the % of individuals that have the allele, Sample size and Location sorted based on their Allele frequency. The results are briefed in the Table 1.

**Table 1.**
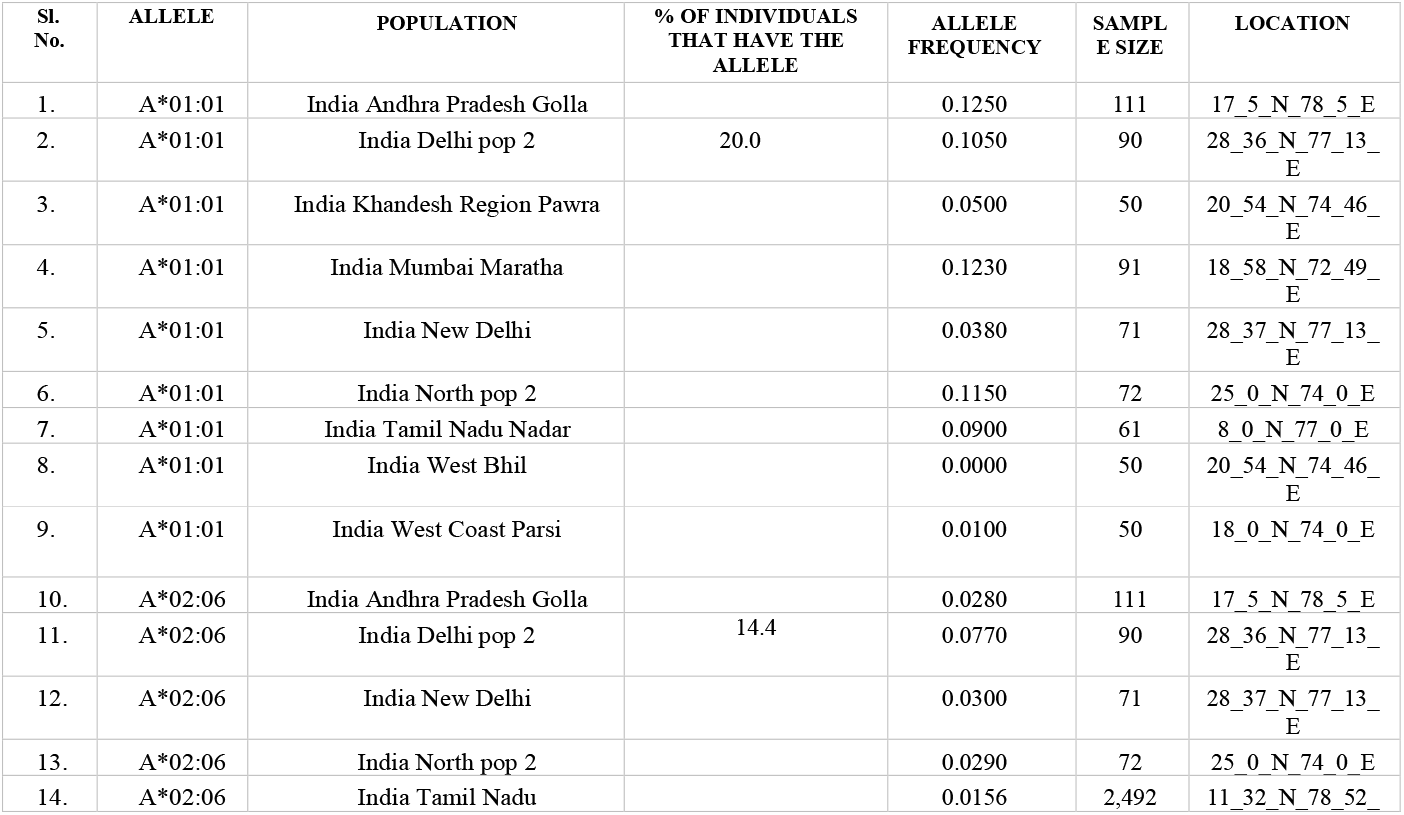
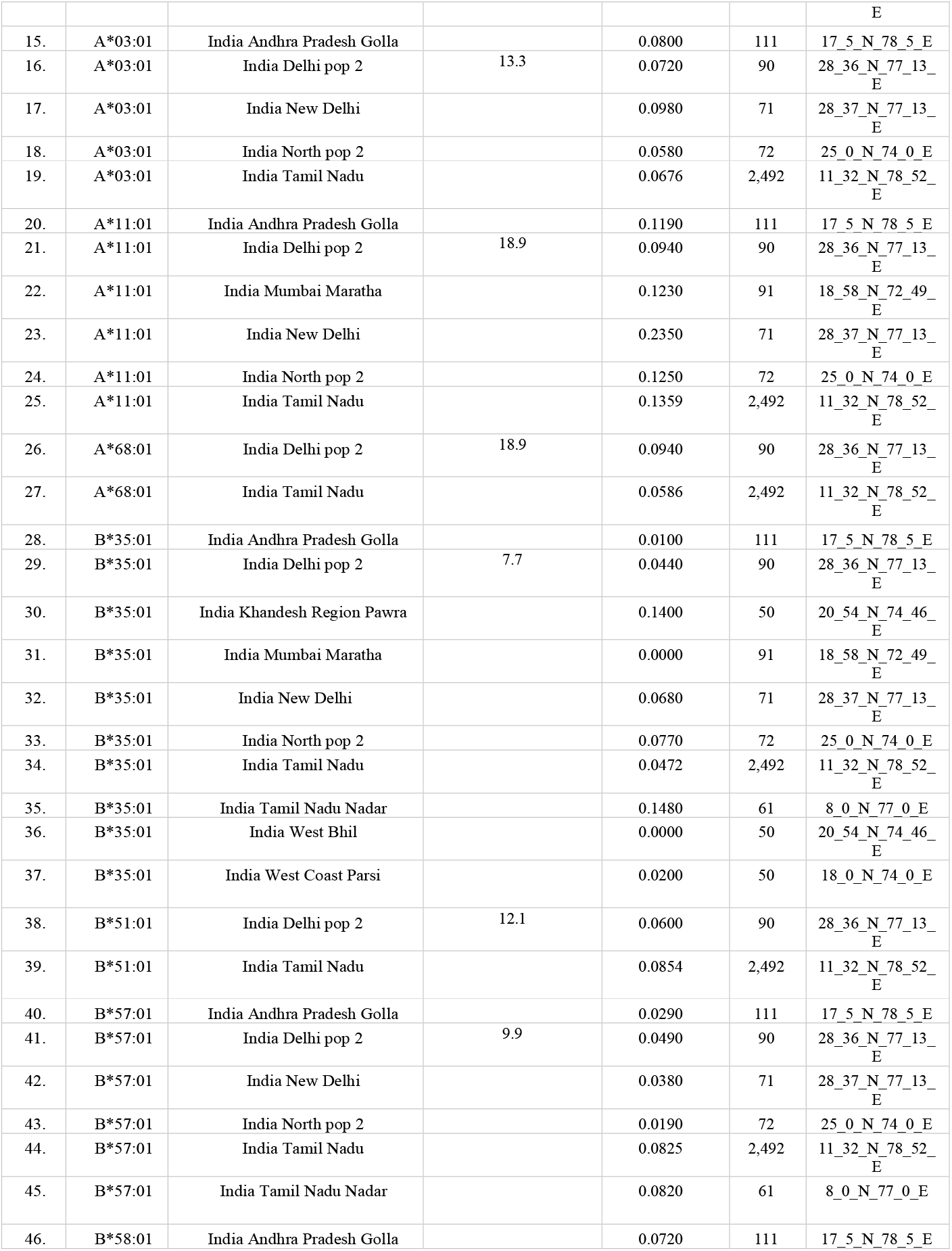
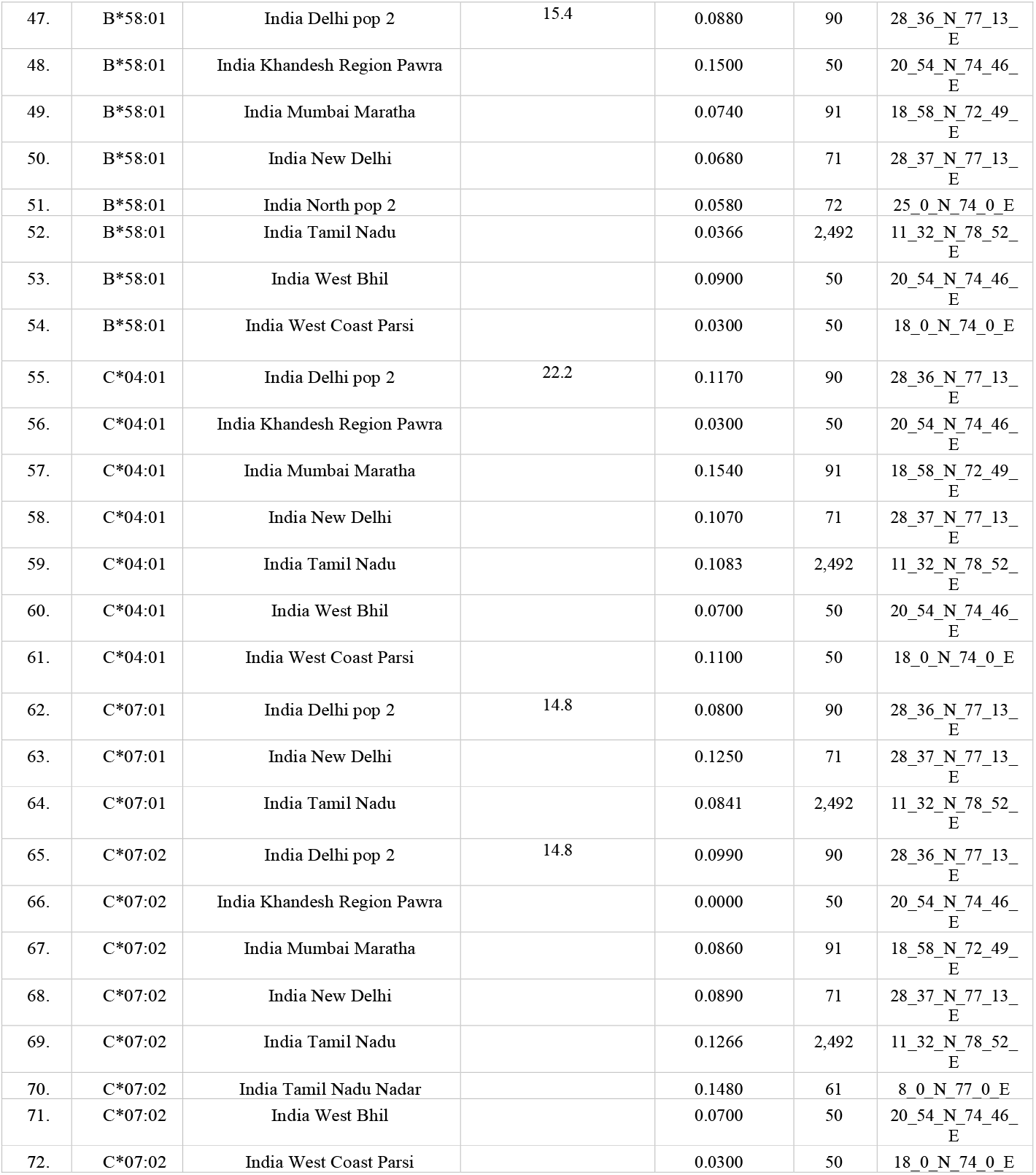
List showing the HLA class I alleles prominent among different regions of Indian population

### 3.6. MHC Class I T-Cell Epitope Prediction

Class-I T cell epitope prediction is done with seven diverse Class-I T-cell epitope prediction tools. These tools make usage of diverse kind of approaches for Class I T-cell prediction which comprises of artificial neural network, PSSM, published motifs etc. Certain tools furthermore predict TAP binding (Epijen v2.0) on the basis of IC50 values and C-terminal of the protein sequence that can undergo proteasomal cleavage (RANKPEP). Range of diverse prediction methods aid to select the best epitopes from the results. Epitopes that are mutual from the results of diverse tools are nominated for further analysis and to evaluate its respond on Indian population using population coverage tool. From the predicted epitopes interacting to different HLA alleles, only those epitopes having decent prediction score and popular among these different tools are selected as it increases the accuracy of the prediction. The results of selected epitopes from all servers are listed in Table S1 under supplementary information.

### 3.7. Population Coverage

T-cell epitopes predicted from 7 different epitope prediction tools are subjected to perform population coverage analysis using IEDB Population Coverage Analysis Tool. This tool basically predicts the fraction of individuals respond to given set of epitopes within the given MHC restriction. 221 selected epitopes common from different prediction servers are given for population coverage analysis. The results have shown the projected population coverage as 83.08% with 19.29 average hit (average number of epitope hits/HLA combinations recognized by the population) and 5.91 pc90 (minimum number of epitope hits/HLA combinations recognized by 90% of the population). A graphical representation of Percent of individuals with Number of epitope hits/HLA combination recognized with cumulative percent of population coverage are obtained as shown in Figure 1 and Table 2. Data of the figure 1 is also presented as tabulated form in supplementray Table S2. Tabulated information on coverage of individual epitopes in Indian population is provided in Table S3 that shows the genotype frequency of various alleles with respect to predicted epitopes.

**Figure 1.**
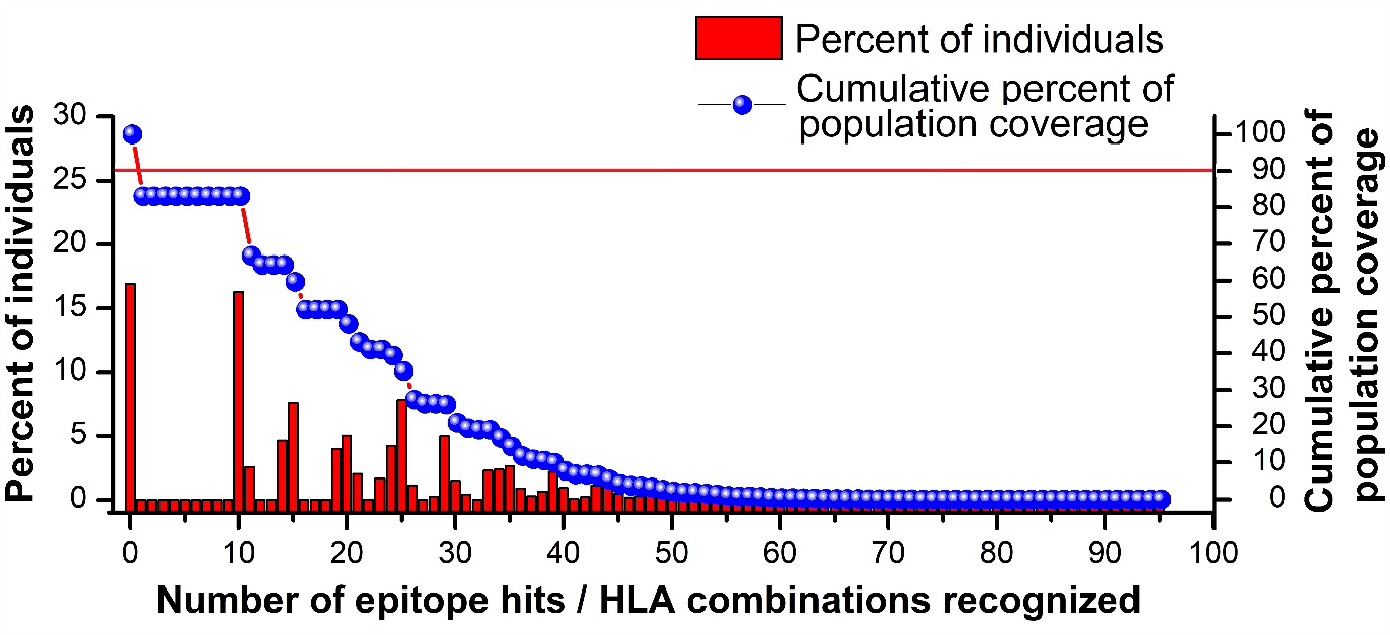
Graphical representation of population coverage analysis reults of MHC-Class I with respect to Indian population.

**Table 2.**
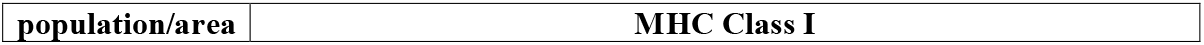
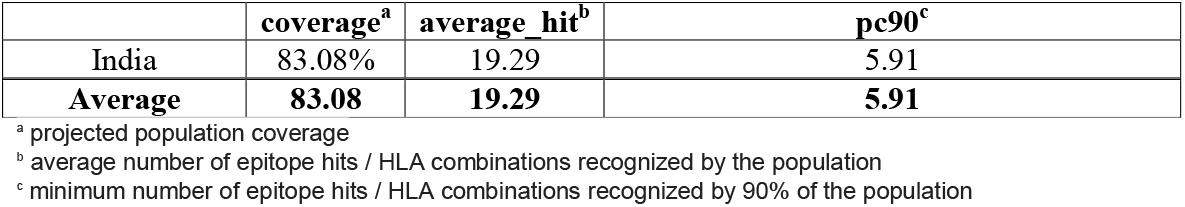
Population Coverage Calculation Result

|  |  |
| --- | --- |
| **population/area** | **MHC Class I** |

|  | **coverage<sup>a</sup>** | **average_hit<sup>b</sup>** | **pc90<sup>c</sup>** |
| --- | --- | --- | --- |
| India | 83.08% | 19.29 | 5.91 |
| **Average** | **83.08** | **19.29** | **5.91** |

<sup>a</sup> projected population coverage
<sup>b</sup> average number of epitope hits / HLA combinations recognized by the population
<sup>c</sup> minimum number of epitope hits / HLA combinations recognized by 90% of the population

### 3.8. Epitope Conservancy and Immunogenicity Prediction

54 epitopes are selected on the basis of their highest coverage among the Indian population. Epitope conservancy prediction using Epitope Conservancy Analysis of IEDB Analysis Resource shows that all the given 54 epitopes are highly conserved within the given spike RBD domain sequence. The percent of protein sequence matches at identity for all the given epitopes are 100% that indicates the high conservancy of the given epitopes. Immunogenicity prediction results from Class I Immunogenicity from IEDB Analysis Resource indicates that most of the given epitopes were immunogenic and capable enough to produce immune response in the human body. As the immunogenic score increases, the immunogenicity of the epitope will be more. The epitope conservancy and immunogenicity results of all the given epitopes are shown in Table 3.

**Table 3.**
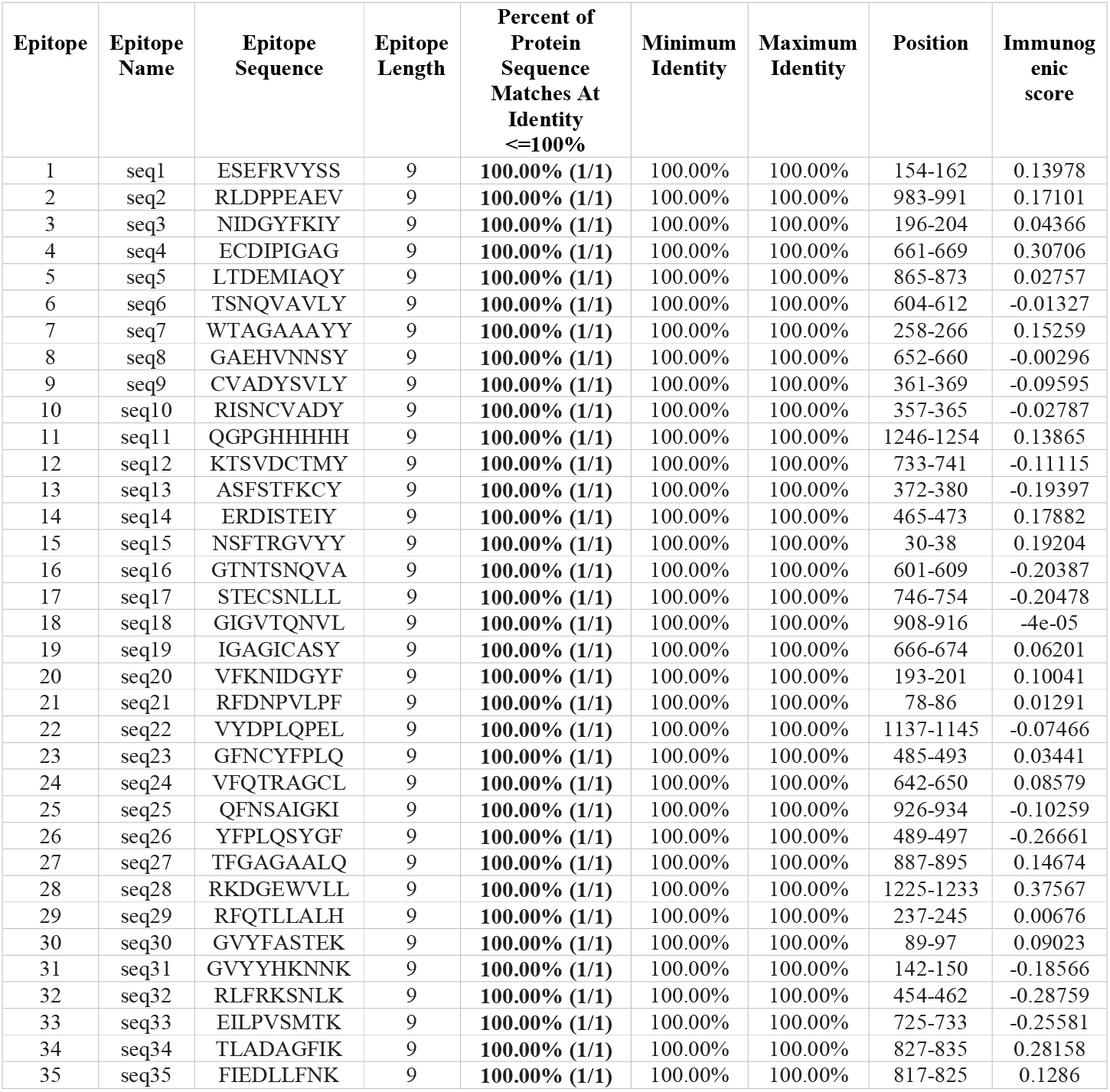
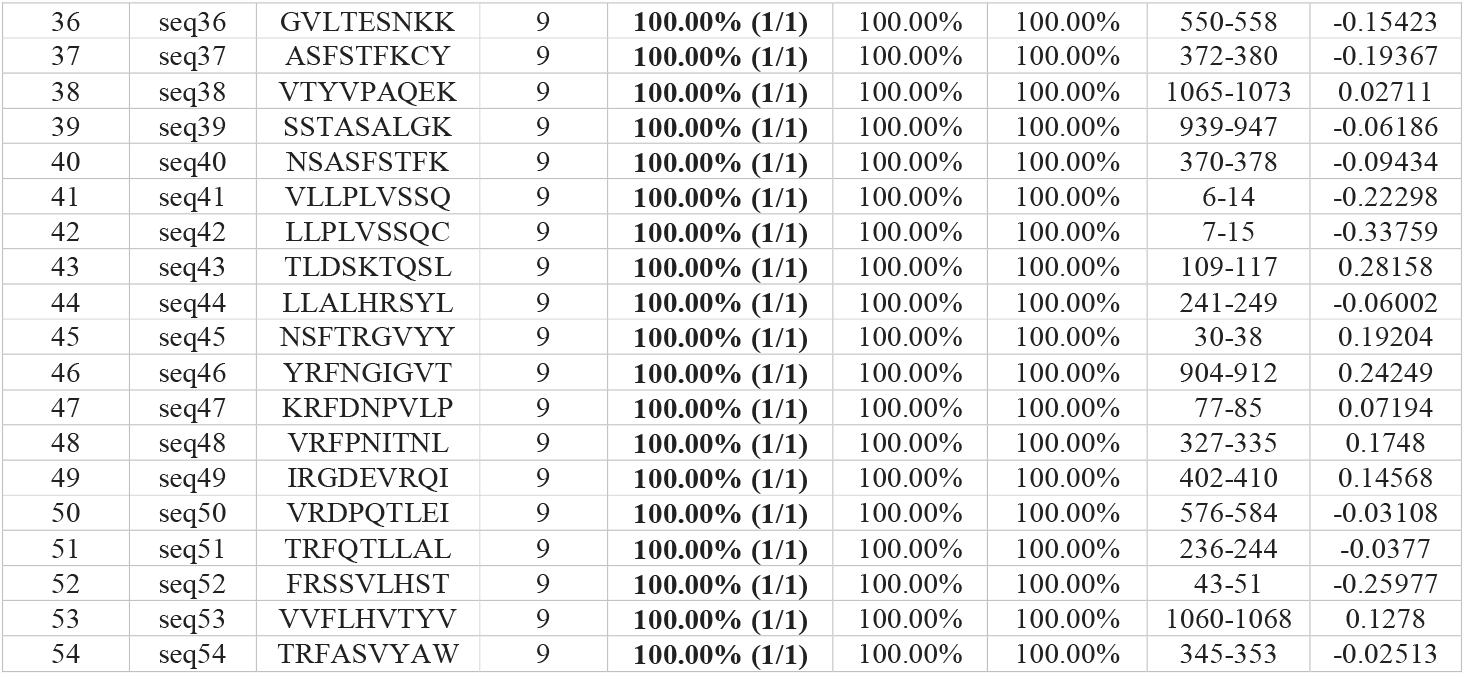
List showing the epitope conservancy and immunogenicity prediction results

### 3.9. Tap Binding Predictions

We have attempted the study of TAP prediction using dataset comprising of 54 9-mer peptides (DS613) that have shown highest population coverage along with high epitope conservancy and immunogenicity and of known affinity to TAP (logIC50relative) using TAPREG. The results are sorted based on increasing TAP affinity IC50(nm) values where one with lowest IC50 values are considered to have highest affinity to TAP proteins. Among the 54 given peptides, these 10 9-mer epitopes having decent IC50 values are chosen to predict their 3D structures for further studies. The results of 10 selected epitopes having high epitope conservancy, immunogenicity score along with decent TAP affinity are summarized in Table 4.

**Table 4.** List showing the tap binding predictions with their IC50 scores of best 5 predicted epitopes:

| NAME | SEQUENCE | TAP AFFINITY<br>IC50 (nM) |
| --- | --- | --- |
| peptide45 | NSFTRGVYY | 39.16 |
| peptide37 | ASFSTFKCY | 47.54 |
| peptide51 | TRFQTLLAL | 114.58 |
| peptide21 | RFDNPVLPF | 234.37 |
| peptide54 | TRFASVYAW | 282.94 |

### 3.10. Allergy Prediction and Toxicity Extrapolation of Epitope

Allergy prediction using AllerTOP v. 2.0 online tool indicate that all the given epitopes are non-allergen in nature. Toxicity prediction is also important to identify whether the predicted epitopes are toxic to human body or not. The allergy and toxicity prediction results of 10 selected epitopes among the 54 given epitopes are shown in Table 5.

**Table 5.** List showing the allergy and toxicity prediction of best 5 epitopes:

| Peptide Sequence | SVM Score | Toxicity Prediction | Allergy Prediction |
| --- | --- | --- | --- |
| NSFTRGVYY | -0.73 | Non-Toxin | Non-Allergen |
| ASFSTFKCY | -0.66 | Non-Toxin | Non-Allergen |
| TRFQTLLAL | -1.04 | Non-Toxin | Non-Allergen |
| RFDNPVLPF | -1.18 | Non-Toxin | Non-Allergen |
| TRFASVYAW | -0.84 | Non-Toxin | Non-Allergen |

### 3.11. Epitope 3-D Structure Prediction

5 non-allergent and non-toxic epitopes are selected on the basis of their TAP binding affinity and immunogenicity. They are NSFTRGVYY, ASFSTFKCY, TRFQTLLAL, RFDNPVLPF and TRFASVYAW. Theses epitopes are submitted to PEP-FOLD server to predict their 3-D structure for further analysis. The model built for each epitope was estimated to analyse their Ramachandran plot using PROCHECK server. It was shown that most portions of the amino acid residues were located in the favoured regions of Ramachandran plot, indicating the high quality of the models for each epitope. It was predicted as around 90.6% (241) residues lie in the most favoured region of the plot whereas 8.6% (23) residues lie in the additional allowable regions. 0.4% (1) residue is found to lie in generously allowed as well as disallowed region of the plot. The 3D structures of epitopes along with the surface structure showing HLA allele-Epitope complex and the Ramachandran plot of the Spike protein sequence is illustrated in figure 2 and figure 3 respectively. Tabulated details of epitope sequence and docking scores are available in table 6.

**Figure 2.**
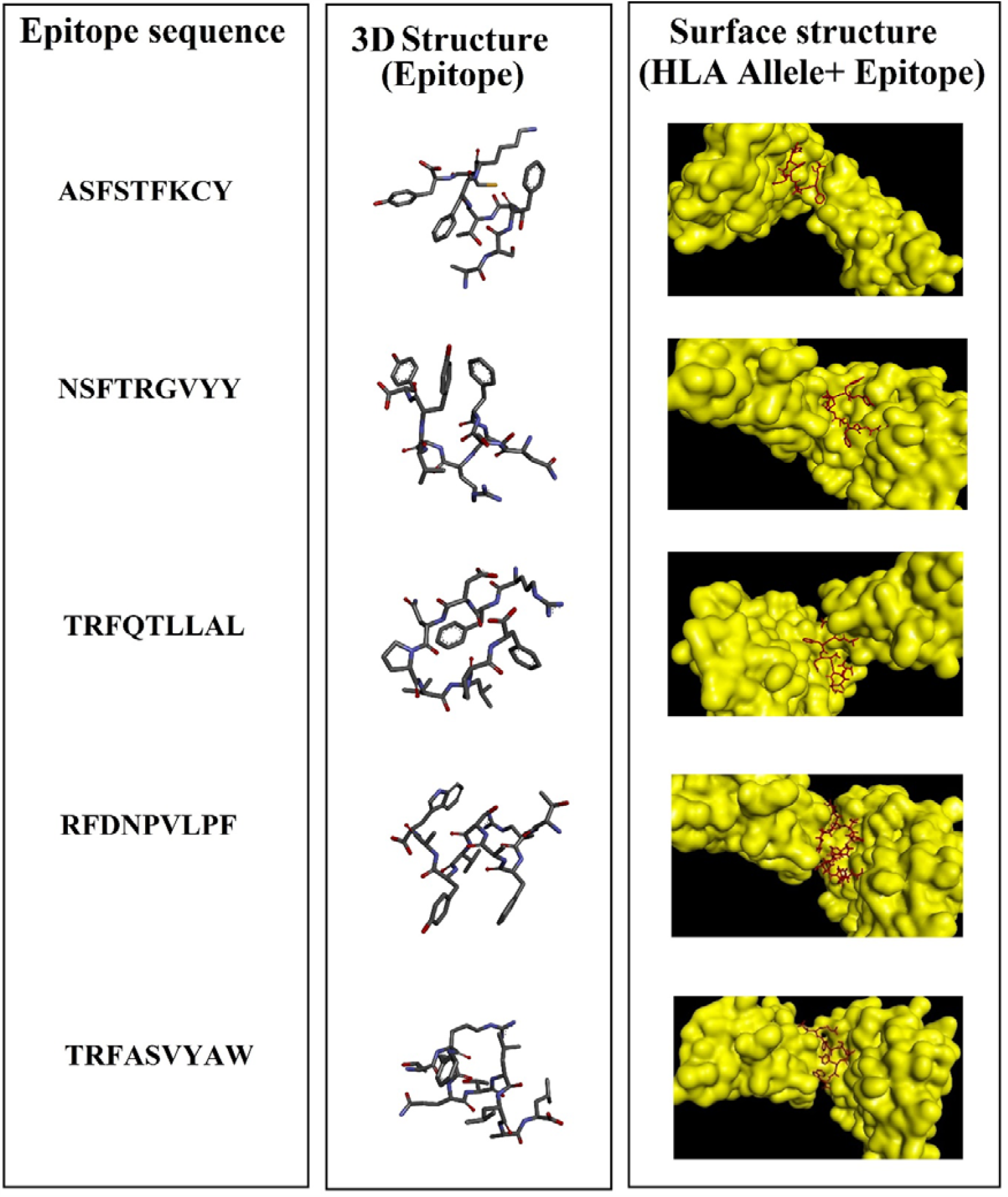
Diagramatic represetnation of epitope sequences seletced on the basis of IC50 score and its 3D structure. Left panel is showing epitope sequences along with its respective 3D strucre and surface structure of complex of epitope and allele in middle and right panel respectively.

**Figure 3.**
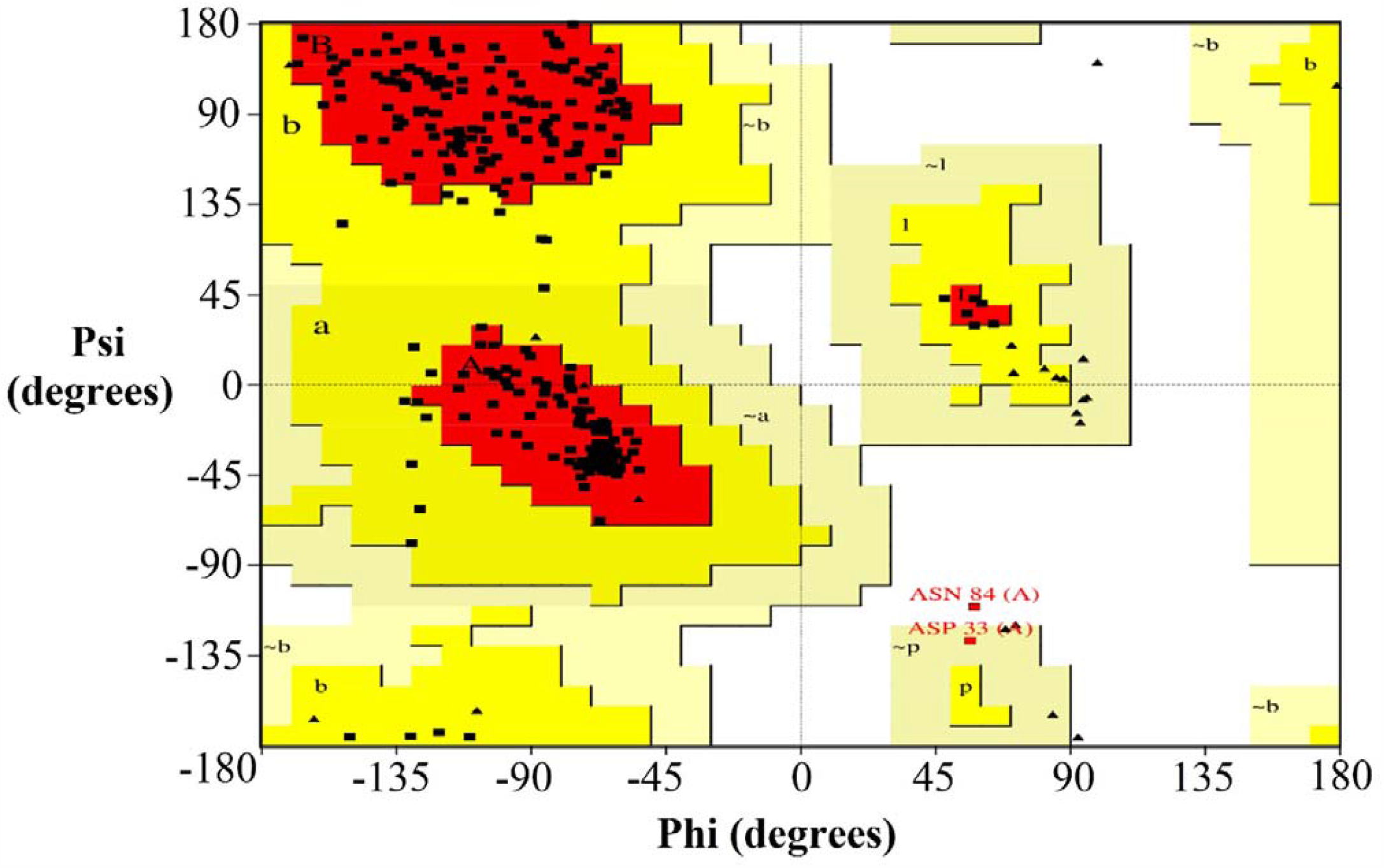
Ramachandran plot of spike protein. It was predicted as around 90.6% (241) residues lie in the most favoured region of the plot whereas 8.6% (23) residues lie in the additional allowable regions. 0.4% (1) residue is found to lie in generously allowed as well as disallowed region of the plot.

**Table 6.**
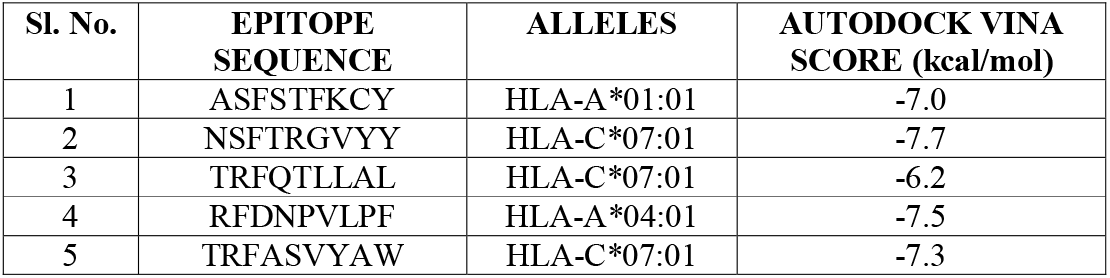
Epitope sequences and allele name and docking score of the Epitope-Allele complex. Docking score were calculated by using Autodock Vina.

### 3.12. Molecular Docking Analysis

Each epitope was docked with their respective alleles to analyse their binding affinities using Autodock Vina. The epitope NSFTRGVYY showed the highest binding affinity with docking score −7.6 kcal/mol with its allele HLA-C*07:01. It was followed by epitopes RFDNPVLPF and TRFASVYAW interacting with alleles HLA-C*04:01 and HLA-C*07:01 respectively. The docked complex was visualized using Pymol and the receptor-ligand interaction was analysed using Discovery Studio. They were having a binding affinity of −7.5 kcal/mol and −7.3 kcal/mol. The docking results and surface structure of 5 epitopes with their respective alleles are illustrated in figure 4.

## 4. DISCUSSION

In the previous few years, many emergent pathogenic diseases have been identified in which most of them have involved zoonotic or species-jumping infectious agents [54]. Among the various emergent viral infections, the novel coronavirus (nCOV or SARS COV 2) is considered as the third CoV outbreak among humans [55]. The COVID-19, emerged in Wuhan, China, at the end of 2019 has been recognized to cause respiratory, digestive, and systematic manifestations that harmfully disturb the human health [56, 57]. This class of virus which affects type 3 pneumocytes and ciliated bronchial epithelial cells using ACE2 receptors are RNA virus and can be transmitted through airborne particles and drops from person to person [58]. As the intensity of these diseases are increasing, vaccine development within a short period is very critical in order to protect the people from the expanding viral attacks. Vaccination is the administration of agent-specific, yet harmless, antigenic particles. Vaccinated individuals can induce defensive immunity against the respective infectious agent when administered. But it would take many years for the progression and production of an effective vaccine and they can be costly too. Therefore, designing of strategies to minimize the cost and time for the development of vaccines become important. In that scenario, various Bioinformatics approaches can be very beneficial to design new-generation safe vaccine within short period of time [59]. By the emergence of technologies like next-generation sequencing, progressive genomics and proteomics, have brought a great transformation in computational immunology [60]. However, with the advancement of new field in Bioinformatics known as the Immunoinformatics which aims in developing the vaccine or vaccine candidates through understanding the immune response of the human body against an organism within a short time. The immunoinformatics is a branch whose core objective is to translate extensive immunological data using computational and mathematical methods, to organize these data to acquire immunologically meaningful elucidations [61 64]. This field uses statistical and machine learning system based tools and can be used for studying and modelling molecular interactions

In the progress of CoV vaccines, numerous approaches are implemented and most of these aim the surface protein named spike (S) glycoprotein or S protein as it is the chief inducer of counteracting antibodies [62]. Spike protein-based approaches of CoV vaccine development either make use of full-length spike protein or S1-receptor-binding domain (RBD) [63]. The spike protein molecule comprises of two subunits namely S1 and S2. The RBD domain present in the S1 subunit interacts with its host cell receptor which is the angiotensin-converting enzyme 2 (ACE2) [64]. Hence, spike protein-based vaccines are considered as vital as it induce antibodies that block not only viral-receptor interaction but also virus genome uncoating. In this proposed immunoinformatics work, we have attempted to recognize class I epitopes from SARS COV2 spike protein with RBD (receptor binding domain). After retrieval of the sequence from NCBI with accession id 6VSB_A, which is the chain A of spike RBD protein, it was exposed to many Insilco approaches to make sure that the epitopes are predicted with higher accuracy. It is vital to identify whether the designated protein sequence is an antigen or not as antigenic or foreign substance can only induce immune response in the host body. The given protein sequence was predicted to be probable antigen with a score of 0.4152. The prediction of physio-chemical parameters of the primary protein sequence help to identify the stability of proteins thereby pointing the stability of epitopes. It also ensures its thermostability, hydrophilicity and theoretical PI. The molecular weight of the protein sequence was computed to be 142274.61, which indicate it to be a good immunogenic sequence with an instability index 31.58 which is appropriate for a stable protein. Three functional domains namely Corona_S2 super family (Coronavirus S2 glycoprotein) of length 601, SARS-CoV-2_Spike_S1_RBD (receptor-binding domain of the S1 subunit of severe acute respiratory syndrome coronavirus 2 Spike (S) protein) of length: 223, Spike-COV-like_S1_NTD (N-terminal domain of the S1 subunit of the Spike (S) protein of length 280 were identified from super secondary prediction. Identification of domains are significant to analyse the functionality as well as to recognize whether the studied protein sequence will be an effective antigen for predicting epitopes. SARS-CoV-2_Spike_S1_RBD is vital as it interacts with the ACE2 receptor in the host. The loop (amino acids 424–494 of the RBD), that comes in whole interaction with the receptor ACE2 is named as receptor-binding motif (RBM) [65]. The role of spike protein especially RBD in receptor binding and membrane fusion show that vaccines targeting spike protein could induce antibodies and T-cell responses to prevent virus binding and fusion or neutralize virus infection. The RBD of S protein comprises of several conformation-dependent epitopes. Epitopes are primarily made by a multi-subunit protease called proteasome who carry out majority of intra-cellular protein degradation [66]. The most accurate C-terminal of CTL epitopes and the N-terminal with a probable extension can be generated by proteosome. If the epitopes are destroyed by the proteasome‘s, CTL responses may diminish. Therefore, to identify possible immunogenic regions in the proteomes of pathogenic microorganisms, the estimation of the proteasome cleavage sites is significant. The spike protein sequence with RBD was predicted to contain 404 cleavage sites within the 1288 amino acid length. So, this specifies that the protein sequence has enough cleavage sites which indicates the presence of decent immunogenic regions that are desirable to generate generous number of epitopes.

Identification of HLA alleles prominent in the Indian population is important to predict the class I T-cell epitopes. Human leukocyte antigen (HLA) loci seemed to be a principal genetic candidate for infectious disease vulnerability [67]. HLAs are categorized as the major histocompatibility complexes (MHCs) due to their significant role in permitting the immune system to distinguish between self and non-self-antigens and are of two types namely class I and class II [68]. Only the alpha chain is variable in class I molecules, so they are named as “HLA”. The three chief class I major histocompatibility complex (MHC) genes are HLA-A, HLA-B, and HLA-C genes. HLA alleles were selected from AFND (Allele Frequency Net Database) on the basis of their allele frequency and percentage of individuals that have the allele. HLA alleles having 9% and above were considered. 12 alleles were selected in which the most prominent one was HLA-C*04:01 (22.2%). The other alleles selected were HLA-A*02:06(14.4%),HLA-A*03:01(13.3%),HLA-A*11:01(18.9%),HLA-A*68:01(18.9%),HLA-B*35:03(13.2%),HLA-B*51:01(12.1%),HLA-B*57:01(9.9%),HLA-B*58:01(15.4%),HLA-A*01:01(20%),HLA-C*07:01(14.8%),HLA-C*07:02(14.8%). The server predicts HLA alleles by covering all regions of India and it also mentions the name of the population in which the alleles are present in higher percentage. Epitopes or antigenic determinant are defined as short amino acid sequences of a protein that can induce a more direct and potent immune response, than the response induced by the whole cognate protein. the strategy for developing epitope vaccines requires an accurate knowledge of the amino acid sequence of the immunogenic protein of interest. They are generally one to six monosaccharides or five to eight −nine amino acid residues on the surface of the antigen. The epitope distinguished by an antibody may reliant upon the occurrence of a specific three-dimensional antigenic conformation known as a conformational epitope. The epitope may also resemble to a simple linear sequence of amino acids and such epitopes are known as linear epitopes. Class I linear T-cell epitopes were predicted using 7 different servers as it can increase the accuracy with of epitope prediction. These servers use different prediction methods including PSSM, neural networks, consensus method etc. to predict 9-mer epitopes or peptides of 9 amino acid length. Epitopes interacting with 12 different class I HLA alleles identified prominently from Indian population were predicted from these servers. The epitopes were selected by comparing results from different servers as it makes sure only exact epitopes are being selected. 212 9-mer epitopes which were mutual in different servers and of decent prediction score were selected for further analysis.

Population coverage calculates the fraction of individuals respond to a given set of epitopes within the given MHC restrictions. For a valuable epitope-based vaccine prediction, population coverage analysis is very significant. It in fact illustrates coverage of all as well as individual epitopes among the population (Supplementary Table S3). As we are interested in Indian population, 212 9-mer epitopes were given as input within class I MHC restrictions. The results show a total coverage of 83.08 with an average hit of 19.29. So, it indicates that among the Indian population, around 83.08% of individuals may respond to the given set of epitopes. Analysis of individual percentage of each epitope showed that highest percentage was 18.85 and were shown by following 10 epitopes VFKNIDGYF, RFDNPVLPF, VYDPLQPEL, GFNCYFPLQ, VFQTRAGCL, QFNSAIGKI, YFPLQSYGF, TFGAGAALQ, RKDGEWVLL and RFQTLLALH. The genotypic frequency of these epitopes was 12.59 which was also highest amongst others and were found to interact with HLA allele HLA-C*04:01. It was tailed by an individual percentage of 18.32% and genotypic frequency of 12.31% shown by 16 epitopes interacting with HLA allele HLA-A*01:01. So, from the population coverage analysis 54 epitopes having highest individual percentage as well as genotypic frequency were chosen. The use of conserved epitopes would be projected to offer wider protection across multiple strains, or even species in an epitope-based vaccine prediction, than epitopes derived from highly variable genome regions. The degree of conservation is defined as the fraction of protein sequences comprising the peptide sequence at a specified identity level. The epitope conservancy analysis of the given peptide sequences showed that the all the epitopes are 100% conserved within the protein sequence. Similarly, immunogenicity prediction of the epitopes is also vital as it is the ability of the epitopes to induce immune response in the host. The immune response to the foreign protein leads to the production of neutralizing antibodies. This immune response is facilitated by T cells and happens as a rapid reaction after its first encounter with the antigen. The prediction of TAP binding affinity of peptides can contribute in subunit vaccine development. The transporter associated with antigen processing (TAP) functions as a transporter of the proteolyzed antigenic or self-altered protein‘s peptide fragments to the endoplasmic reticulum where the binding of these peptides with the major histocompatibility complex (MHC) class I molecules occurs [69]. Consequently, prediction of TAP-binding peptides is extremely beneficial in recognizing the MHC class I-restricted T-cell epitopes. TAP affinity which is measured in terms of IC50 score as lower the IC50 score more will be the binding affinity of epitopes with the TAP proteins. Among the given 54 epitopes, the highest affinity was shown by the sequence NSFTRGVYY with an IC50 score of 39.16 nM followed by ASFSTFKCY with 47.54 nM. Then these 54 epitopes were analysed for their allergy and toxicity prediction. Allergy prediction of the epitopes is vital to detect whether the epitopes can trigger IgE antibodies in the human host. From allergy prediction it was found that all the epitopes are non-allergen and appropriate for developing T-cell based epitope vaccines. Similarly, toxicity prediction also showed that all the given epitopes are non-toxic to the host. Based on the outcomes from epitope conservancy analysis, immunogenicity prediction, TAP binding prediction allergy and toxicity prediction, 5 succeeding non-allergen and non-toxic epitopes NSFTRGVYY, ASFSTFKCY, TRFQTLLAL, RFDNPVLPF and TRFASVYAW having appropriate immunogenicity score, epitope conservancy and IC50 score were favoured for additional analysis. All these were having 100% epitope conservancy with decent immunogenicity score and IC50 values within 2500nM.

The 5 selected epitopes NSFTRGVYY, ASFSTFKCY, TRFQTLLAL, RFDNPVLPF and TRFASVYAW were given for their 3D structure prediction. 3-D structure prediction of epitopes is necessary to perform molecular docking and other structural analysis. The predicted structure was downloaded in PDB format and were found to lie in the favourable region of the Ramachandran Plot. Molecular docking analysis of the epitopes with their respective HLA alleles shown that all the epitopes are interacting with their allele with a decent binding affinity value. The highest binding score was shown by NSFTRGVYY (−7.6 kcal/mol) followed by RFDNPVLPF (−7.5 kcal/mol) and TRFASVYAW (−7.3 kcal/mol). The output pdbqt file having different ligand pose was split using vina split to choose the one with best pose. The docked complex was visualized in Pymol. The interactions as well as the surfaces were analysed using Discovery Studio tool. One of the epitopes TRFASVYAW among 3 best interacting epitopes is seen within the RBD domain (sequence 345-353) of spike glycoprotein which indicates that it might be considered as a major epitope for vaccine design to inhibit viral attachment with the host receptor.

## 5. CONCLUSION

The SARS-CoV-2 virus has become a key health concern in many countries including India and have led to death of many people worldwide. Although the development of vaccine is on progress but in this difficult situation, treatment only depend on the antiretroviral therapy [70]. Therefore, this study might provide an add-on to the path of multi-epitope-based vaccine development against SARS-CoV-2. Among different proteins of SARS-CoV-2, we have chosen RBD domain of spike glycoprotein as it is crucial for viral attachment with host receptor in FASTA format. The conserved sequence of spike glycoprotein with RBD domain of the infectious SARS-CoV-2 was evaluated using the advanced immunoinformatics methods.

Immunoinformatics can effectively make use of computational techniques to bring out effective and useful advantage in the exploration of novel vaccines [71]. It is believed to contribute to vaccine design as the computational chemistry contributes to drug design. Immunoinformatics-based vaccine design can attain worthy and cost-efficient advancement in vaccines or vaccine components design. It has been reported that the CTL epitope-based vaccine, which has an elevated realism in vaccine designing can complement the convalescent plasma therapy and can produce multiple serotype-specifc immune responses [72]. The present study concludes that the highly potential epitope vaccine construct has the potential to obtain sturdy immune responses. The designed epitopes were assessed over several immunological parameters. The epitopes exhibited high antigenicity, immunogenicity, and C-terminal proteasomal affinity and strong TAP affinity. They were also identified as non-allergen, highly antigenic along with stable physio-chemical characterization. At last, molecular docking and dynamics study was performed to check the binding affinity of predicted epitopes with its respective HLA alleles.

Current immunoinformatic analysis pointed out best 5 MHC □ I epitopes within the spike glycoprotein RBD of SARS□COV□2 that can be used for designing multi-epitope vaccine Indian population. The population coverage analysis given epitopes within Indian population has helped to identify best 5 epitopes that responds more with the population and might provide an insight for multi-epitope-based vaccine as the disease has caused a major impact on the country. Designing a vaccine specifically for a population is helpful as different population may have different response to the epitopes and as a result a general vaccine may not give the expected results. Yet, these immunoinformatic analyses need several in vitro and in vivo validations before formulating the vaccine to resist COVID□19. In-silico studies of the predicted epitopes portrayed decent elicitation of anti-corona immune response. This study might provide a way to a potential CTL based vaccine construct against SARS-CoV-2 through detailed theoretical analysis.

## 6. Declaration

### 6.2. Consent to participate

All authors has given their consent to participate in this work.

### 6.3. Consent for publication

All authors has given their consent to publish this work.

### 6.4. Availability of data and material

Not applicable.

### 6.5. Code availability

Not applicable.

### 6.6. Authors‘ contributions

MD has conceptualized and supervised the work. SD has performed the experiments. SD and MD have analysed the data and written the manuscript.

## 6.7. Acknowledgement

MD thanks to the DST-INSPIRE Faculty award (2017), Govt. of India.

## REFERENCES

1. Mackay, Ian R.; Rosen, Fred S.; Delves, Peter J.; Roitt, Ivan M. (2000) The Immune System. New England Journal of Medicine, 343(1),37–49.

2. Korber, Bette; LaBute, Montiago; Yusim Karina (2006) Immunoinformatics Comes of Age. PLoS Computational Biology, 2(6), e71.

3. Hanada K, Yewdell JW, Yang JC (2004) Immune recognition of a human renal cancer antigen through post-translational protein splicing. Nature 427: 252–256.

4. Janeway CA, Travers P, Walport M, Shlomchik M (2005) Immunobiology. New York: Garland Science Publishing. 600 p.

5. Rudolph MG, Stanfield RL, Wilson IA (2006) How TCRs bind MHCs, peptides, and coreceptors. Annu Rev Immunol 24: 419–466.

6. Maenaka K, Jones EY (1999) MHC superfamily structure and the immune system. Curr Opin Struct Biol 9: 745–753.

7. Messaoudi I, LeMaoult J, Metzner BM, Miley MJ, Fremont DH, et al. (2001) Functional evidence that conserved TCR CDR alpha 3 loop docking governs the cross-recognition of closely related peptide:class I complexes. J Immunol 167: 836–843.

8. Saphire EO, Parren PW, Pantophlet R, Zwick MB, Morris GM, et al. (2001) Crystal structure of a neutralizing human IGG against HIV-1: A template for vaccine design. Science 293: 1155–1159.

9. Tomar N, De RK. (2010) Immunoinformatics: an integrated scenario. Immunology. 131(2):153–68.

10. Doytchinova, I. A., & Flower, D. R. (2007) VaxiJen: a server for prediction of protective antigens, tumour antigens and subunit vaccines. BMC Bioinformatics, 8(1), 4.

11. Davies MN, Flower DR (2007) Harnessing bioinformatics to discover new vaccine. Drug Discovery Today; 12(9-10) 389–95.

12. Bhattacharya M, Sharma AR, Patra P, Ghosh P, Sharma G, Patra BC, Lee SS, Chakraborty C. (2020) Development of epitope-based peptide vaccine against novel coronavirus 2019 (SARS-COV-2): Immunoinformatics approach. J Med Virol. 92(6):618–631.

13. Zhou P, Yang XL, Wang XG, Hu B, Zhang L, Zhang W, Si HR, Zhu Y, Li B, Huang CL, Chen HD, Chen J, Luo Y, Guo H, Jiang RD, Liu MQ, Chen Y, Shen XR, Wang X, Zheng XS, Zhao K, Chen QJ, Deng F, Liu LL, Yan B, Zhan FX, Wang YY, Xiao GF, Shi ZL. (2020) A pneumonia outbreak associated with a new coronavirus of probable bat origin. Nature. 579(7798):270–273.

14. Zhu N, Zhang D, Wang W, Li X, Yang B, Song J, Zhao X, Huang B, Shi W, Lu R, Niu P, Zhan F, Ma X, Wang D, Xu W, Wu G, Gao G-F, Tan W, (2020). A novel coronavirus from patients with pneumonia in China, 2019. N Engl J Med. 2020; 382:727□733

15. Chan JF□W, Yuan S, Kok K□H, Wang K-K, Chu H, Yang J, Xing F, Liu J, Yip C-Y, Wing R, Shan R-W, Tsoi W-H, Lo S K-F, Chan K-H, Man V-K, Chan W-M, Daniel J, Cai J-P, Cheng V-C, Yuen K-Y (2020). A familial cluster of pneumonia associated with the 2019 novel coronavirus indicating person□to□person transmission: a study of a family cluster. The Lancet; 395:514□523.

16. Carlos WG, Dela Cruz CS, Cao B, Pasnick S, Jamil S (2020). Novel Wuhan (2019□nCoV) coronavirus. Am J Respir Crit Care Med. 201: P7□P8.

17. Perlman S (2020). Another decade, another coronavirus. N Engl J Med.; 382(8):760–762.

18. Yin D, Li L, Song X, Li Han, Wang J, Ju W, Qu X, Song D, Liu Y, Meng X, Cao H, Song W, Meng R, Liu J, Li Juan, Xu K (2016). A novel multi-epitope recombined protein for diagnosis of human brucellosis. BMC Infect Dis;16(1):219.

19. Chung M, Bernheim A, Mei X, Zhang N, Huang M, Zeng X, Cui J, Xu W, Yang Y, Fayad ZA, Jacobi A, Li K, Li S, Shan H. (2020) CT Imaging Features of 2019 Novel Coronavirus (2019-nCoV). Radiology. 295(1):202–207.

20. NCBI Resource Coordinators. (2018) Database resources of the National Center for Biotechnology Information. Nucleic Acids Res. 4;46(D1):D8–D13.

21. Doytchinova, I. A., & Flower, D. R. (2007). Identifying candidate subunit vaccines using an alignment-independent method based on principal amino acid properties. Vaccine, 25(5), 856–866.

22. Gasteiger E., Hoogland C., Gattiker A., Duvaud S., Wilkins M.R., Appel R.D., Bairoch A (2005); Protein Identification and Analysis Tools on the ExPASy Server; (In) John M. Walker (ed): The Proteomics Protocols Handbook, Humana Press. pp. 571–607

23. Gill, S.C. and von Hippel, P.H. (1989) Calculation of protein extinction coefficients from amino acid sequence data. Anal. Biochem. 182:319–326

24. Bachmair, A., Finley, D. and Varshavsky, A. (1986) In vivo half-life of a protein is a function of its amino-terminal residue. Science 234, 179–186

25. Guruprasad, K., Reddy, B.V.B. and Pandit, M.W. (1990) Correlation between stability of a protein and its dipeptide composition: a novel approach for predicting in vivo stability of a protein from its primary sequence. Protein Eng. 4,155–161.

26. Ikai, A.J. (1980) Thermostability and aliphatic index of globular proteins. J. Biochem. 88, 1895–1898.

27. Kyte, J. and Doolittle, R.F. (1982) A simple method for displaying the hydropathic character of a protein. J. Mol. Biol. 157, 105–132

28. Lu S, Wang J, Chitsaz F, Derbyshire MK, Geer RC, Gonzales NR, Gwadz M, Hurwitz DI, Marchler GH, Song JS, Thanki N, Yamashita RA, Yang M, Zhang D, Zheng C, Lanczycki CJ, Marchler-Bauer A. (2020) CDD/SPARCLE: the conserved domain database in 2020. Nucleic Acids Res. 8;48(D1):D265–D268.

29. El-Gebali, S., Mistry, J., Bateman, A., Eddy, S.R., Luciani, A., Potter, S.C., Qureshi, M., Richardson, L.J., Salazar, G.A., Smart, A. et al. (2019) The Pfam protein families database in 2019. Nucleic Acids Res., 4D427–D432.

30. Letunic, I. and Bork, P. (2018) 20 years of the SMART protein domain annotation resource. Nucleic Acids Res., 46, D493–D496.

31. Tatusov, R.L., Natale, D.A., Garkavtsev, I.V., Tatusova, T.A., Shankavaram, U.T., Rao, B.S., Kiryutin, B., Galperin, M.Y., Fedorova, N.D. and Koonin, E.V. (2001) The COG database: new developments in phylogenetic classification of proteins from complete genomes. Nucleic Acids Res., 29, 22–28.

32. Haft, D.H., Selengut, J.D., Richter, A.R., Harkins, D., Basu, M.K. and Beck, E. (2013) TIGRFAMs and genome properties in 2013. Nucleic Acids Res., 41, D387–D395.

33. M. Nielsen, C. Lundegaard, O. Lund, and C. Kesmir(2005) The role of the proteasome in generating cytotoxic T cell epitopes: Insights obtained from improved predictions of proteasomal cleavage. Immunogenetics., 57(1-2):33–41, 2005.

34. Gonzalez-Galarza FF, McCabe A, Santos EJMD, Jones J, Takeshita L, Ortega-Rivera ND, Cid-Pavon GMD, Ramsbottom K, Ghattaoraya G, Alfirevic A, Middleton D, Jones AR. (2020) Allele frequency net database (AFND) 2020 update: gold-standard data classification, open access genotype data and new query tools. Nucleic Acids Res. 8;48(D1):D783–D788.

35. Reche PA, Glutting JP, Zhang H, Reinherz EL (2004). Enhancement to the RANKPEP resource for the prediction of peptide binding to MHC molecules using profiles. 456:405–419

36. Hans-Georg Rammensee, Jutta Bachmann, Niels Nikolaus Emmerich, Oskar Alexander Bachor, Stefan Stevanovic (1999): SYFPEITHI: database for MHC ligands and peptide motifs. Immunogenetics 50: 213–219

37. Ayyoub, M., Stevanovic, S., Sahin, U., Guillaume, P., Servis, C., Rimoldi, D., Valmori, D., Romero, P., Cerottini, J.C., Rammensee, HG., Pfreundschuh, M., Speiser, D. and Levy, F. (2002) Proteasome-assisted identification of a SSX-2-derived epitope recognized by tumor-reactive CTL infiltrating metastatic melanoma. J. Immunol., 168, 1717–22.

38. Brusic V., Rudy G., Honeyman M.C., Hammer J. and Harrison L.C. (1998). Prediction of MHC class-II binding peptides using an evolutionary algorithm and artificial neural network. Bioinformatics 14(2), 121–130.

39. Vita R, Mahajan S, Overton JA, Dhanda SK, Martini S, Cantrell JR, Wheeler DK, Sette A, Peters B (2018). The Immune Epitope Database (IEDB): 2018 update. Nucleic Acids Res. 2018 Oct 24.

40. Doytchinova, I. A., P. Guan, D. R. Flower (2006). EpiJen: a server for multi-step T cell epitope prediction. BMC Bioinformatics, 7, 131

41. Jurtz V, Paul S, Andreatta M, Marcatili P, Peters B, Nielsen M. NetMHCpan-4.0: Improved Peptide-MHC Class I Interaction Predictions Integrating Eluted Ligand and Peptide Binding Affinity Data. J Immunol. 2017 Nov 1;199(9):3360–3368.

42. Bui H. H, Sidney J, Dinh K, Southwood S, Newman M. J, Sette A. (2006). Predicting population coverage of T-cell epitope-based diagnostics and vaccines. BMC Bioinformatics 17:153.

43. Bui H. H,Sidney J, Li W, Fusseder N, Sette A., (2007). Development of an epitope conservancy analysis tool to facilitate the design of epitope-based diagnostics and vaccines. BMC Bioinformatics 8(1):361

44. Calis JJA, Maybeno M, Greenbaum JA, Weiskopf D, De Silva AD, Sette A, Kesmir C, Peters B. (2013). Properties of MHC class I presented peptides that enhance immunogenicity. PloS Comp. Biol. 8(1):361.

45. Diez-Rivero CM, Chenlo B, Zuluzga P and Reche PA (2010) Quantitative modeling of peptide binding to TAP using support vector machine. Proteins, 78:63–72

46. Dimitrov, I., Bangov, I., Flower, D. R., & Doytchinova, I. (2014) AllerTOP v.2—a server for in silico prediction of allergens. Journal of Molecular Modeling 20(6).

47. Gupta S, Kapoor P, Chaudhary K, Gautham, A., Kumar, R., Raghava, G.P.S (2013) In silico Approach for Predicting Toxicity of Peptides and Proteins. PLoS ONE 8(9): e73957

48. Beaufays J, Lins L, Thomas A, Brasseur R. (2012) In silico predictions of 3D structures of linear and cyclic peptides with natural and non-proteinogenic residues. J Pept Sci. 18(1):17–24.

49. Levine MM, Lagos R, Esparza J (2016). Vaccines and vaccination in historical perspective. In New Generation Vaccines. 2nd edition 1–11.

50. Ada GL (1997). The traditional vaccines: an overview. In New Generation Vaccines. 2nd edition 13–23.

51. Davies MN, Flower DR (2007). Harnessing bioinformatics to discover new vaccine. Drug Discov Today; 12:389–95.

52. Laskowski R A, MacArthur M W, Moss D S, Thornton J M (1993). PROCHECK - a program to check the stereochemical quality of protein structures. J. App. Cryst., 26, 283–291.

53. Trott O, Olson AJ. (2010) AutoDock Vina: improving the speed and accuracy of docking with a new scoring function, efficient optimization, and multithreading. J Comput Chem. 30;31(2):455–61.

54. D. Katterine Bonilla-Aldana1,2, Kuldeep Dhama3, Alfonso J. Rodriguez-Morales (2020). Revisiting the One Health Approach in the Context of COVID-19: A Look into the Ecology of this Emerging Disease. Adv. in Ani. and Vet. Sci. 8 (3) 234–237

55. Munster VJ, Koopmans M, van Doremalen N, van Riel D, de Wit E. (2020) A Novel Coronavirus Emerging in China - Key Questions for Impact Assessment. N Engl J Med. 20;382(8):692–694.

56. Li Q, Guan X, Wu P, Wang X, Zhou L, Tong Y, Feng Z (2020). Early Transmission Dynamics in Wuhan, China, of Novel Coronavirus-Infected Pneumonia. N. Engl. J. Med.

57. Liu SL, Saif L (2020). Emerging Viruses without Borders: The Wuhan Coronavirus. Viruses. 12(2).

58. Rodriguez-Morales AJ, MacGregor K, Kanagarajah S, Patel D, Schlagenhauf P (2020). Going global - Travel and the 2019 novel coronavirus. Travel Med. Infect. Dis. 33: 101578.

59. María, Ribas□Aparicio & Castelán, Juan & Alicia, Jiménez□ & Paulina, Monterrubio□López & Gerardo, Aparicio□. (2017). The Impact of Bioinformatics on Vaccine Design and Development. Intechopen, chap 7 (DOI:10.5772/INTECHOPEN.69273).

60. Backert, L., Kohlbacher, O. Immunoinformatics and epitope prediction in the age of genomic medicine. Genome Med 7, 119.

61. Patronov A, Doytchinova I. T-cell epitope vaccine design by immunoinformatics. Open Biol. 2013 Jan 8;3(1):120139.

62. Du, L., He, Y., Zhou, Y., Liu, S., Zheng, B.-J., & Jiang, S. (2009). The spike protein of SARS-CoV — a target for vaccine and therapeutic development. Nature Reviews Microbiology, 7(3), 226–236.

63. Wang, Q., Wong, G., Lu, G., Yan, J., & Gao, G. F. (2016). MERS-CoV spike protein: Targets for vaccines and therapeutics. Antiviral Research, 133, 165–177.

64. Li, F. (2005). Structure of SARS Coronavirus Spike Receptor-Binding Domain Complexed with Receptor. Science, 309(5742), 1864–1868.

65. Wong, S. K., Li, W., Moore, M. J., Choe, H. & Farzan, M (2004). A 193-amino acid fragment of the SARS coronavirus S protein efficiently binds angiotensin-converting enzyme 2. J. Biol. Chem. 279, 3197–3201

66. Nielsen, M., Lundegaard, C., Lund, O., & Keşmir, C. (2005). The role of the proteasome in generating cytotoxic T-cell epitopes: insights obtained from improved predictions of proteasomal cleavage. Immunogenetics, 57(1-2), 33–41.

67. Nguyen A, David JK, Maden SK, et al. (2020) Human Leukocyte Antigen Susceptibility Map for Severe Acute Respiratory Syndrome Coronavirus 2. J Virol. 94(13):e00510–20.

68. Crux, N. B., & Elahi, S. (2017). Human Leukocyte Antigen (HLA) and Immune Regulation: How Do Classical and Non-Classical HLA Alleles Modulate Immune Response to Human Immunodeficiency Virus and Hepatitis C Virus Infections? Front Immunol. 2017 Jul 18;8:832.

69. Bhasin, M. (2004). Analysis and prediction of affinity of TAP binding peptides using cascade SVM. Protein Science, 13(3), 596–607.

70. Kumar, Neeraj & Admane, Nikita & Kumari, Anchala & Sood, Damini & Grover, Sonam & Prajapati, Vijay & Chandra, Ramesh & Grover, Abhinav. (2020). Cytotoxic T-Lymphocyte Elicited Vaccine against SARS-CoV-2 employing Immunoinformatics Framework. Research square preprint server (DOI: https://doi.org/10.21203/rs.3.rs-40659/v1).

71. Bhattacharya M, Sharma AR, Patra P, Ghosh P, Sharma G, Patra BC, Lee SS, Chakraborty C. Development of epitope-based peptide vaccine against novel coronavirus 2019 (SARS-COV-2): Immunoinformatics approach. J Med Virol. 2020 Jun;92(6):618–631.

72. Pandey, R. K., Ojha, R., Aathmanathan, V. S., Krishnan, M., & Prajapati, V. K. (2018). Immunoinformatics approaches to design a novel multi-epitope subunit vaccine against HIV infection. Vaccine, 36(17), 2262–2272.

