## Supplementary Table S1, S2, S3 for "Immunoinformatic based analytics on T-cell epitope from spike protein of SARS-CoV-2 concerning Indian population"

**Supplementary Information**

**Table S1.** List showing epitopes predicted to interact with given HLA alleles from 7 different epitope prediction servers along with the sequence position of epitopes

| SERVER | ALLELE | SEQUENCE | POSITION |
| --- | --- | --- | --- |
| 1)RANKPEP | HLA-A*01:01  HLA-A*02:06  HLA-A*03:01  HLA-B*07:02  HLA-B*35:01  HLA-B*51:01  HLA-A*29:02  HLA-A*11:01  HLA-A*68:01  HLA-B*58:01  HLA-B*57:01 | ESEFRVYSS  RLDPPEAEV  NIDGYFKIY  QGPGHHHHH  ECDIPIGAG  LTDEMIAQY  KTSVDCTMY  ERDISTELY  YLQPRTFLL  ELLHAPATV  FCNDPFLGV  KIADYNYKL  RLFRKSNLK  GIYQTSNFR  GVYYHKNNK  EILPVSMTK  QIYKTPPIK  IPFAMQMAY  FPQSAPHGV  IPTNFTISV  QPTESIVRF  APRDGQAYV  LPFNDGVYF  IPIGAGICA  LPPAYTNSF  CPFGEVFNA  LPDDFTGCV  LPPLLTDEM  LPFFSNVTW  IPFAMQMAY  TPINLVRDL  TPCNGVEGF  LPPAYTNSF  FAMQMAYRF  DPPEAEVQI  IAIPTNFTI  LPPLLTDEM  LPPAYTNSF  LPLVSSQCV  PPLLTDEMI  IAPGQTGKI  FPQSAPHGV  LPFNDGVYF  LPFFSNVTW  GEVFNATRF  QEVFAQVKQ  AEIRASANL  FERDISTEI  SKVGGNYNY  GVYFASTEK  GVYYHKNNK  RLFRKSNLK  EILPVSMTK  TLADAGFIK  FIEDLLFNK  GVLTESNKK  ASFSTFKCY  VTYVPAQEK  GVYFASTEK  GTHWFVTQR  EILPVSMTK  RLFRKSNLK  SVYAWNRKR  GVLTESNKK  HADQLTPTW  LTDEMIAQY  LAGTITSGW  NATNVVIKV  SGTNGTKRF  QPTESIVRF  DFTGCVIAW  VASQSIIAY  FAMQMAYRF  KTPPIKDFG  FTNVYADSF  QSAPHGVVF  HADQLTPTW  DFTGCVIAW  HHHHHHSAW  LAGTITSGW  TSNQVAVLY  KSFTVEKGI | 154  983  196  1246  661  865  733  465  269  516  135  417  454  311  142  725  787  896  1052  714  321  1216  84  664  24  336  425  861  56  896  208  478  24  898  985  712  861  24  8  862  410  1052  84  56  339  779  1016  464  443  89  142  454  725  827  817  550  372  1065  89  1099  725  454  349  550  625  865  878  122  71  321  428  687  898  790  392  1054  625  428  1252  878  604  304 |
| 2)SYFPEITHI | HLA-A*11:01  HLA-A*68:01  HLA-B*07:02  HLA-B*50:01 | SSTASALGK  GVYFASTEK  NSASFSTFK  VTYVPAQEK  GTHWFVTQR  SVYAWNRKR  QIAPGQTGK  IADTTDAVR  NASVVNIQK  FVIRGDEVR  NSASFSTFK  GVLTESNKK  APRDGQAYV  TPINLVRDL  APGQTGKIA  IPIGAGICA  LPFNDGVYF  CPFGEVFNA  QPTESIVRF  LPDDFTGCV  GPKKSTNLV  SPGSASSVA  IPTNFTISV  KPFERDIST  SEFRVYSSA  TEVPVAIHA  NENGTITDA  YECDIPIGA  PEAPRDGQA  FELLHAPAT  RAAEIRASA  TTRTQLPPA  MESEFRVYS  FEYVSQPFL  RDLPQGFSA | 939  89  370  1065  1099  349  409  569  1173  400  370  550  1216  208  411  664  84  336  321  425  526  680  714  462  155  618  280  660  1214  515  1014  19  153  168  214 |
| 3)ANNPRED | HLA-A*02:06  HLA-A*03:01  HLA-A*11:01 | LLHAPATVC  VLSFELLHA  GVGYQPYRV  VLLPLVSSQ  LLPLVSSQC  YLQPRTFLL  SWMESEFRV  TLDSKTQSL  IVNNATNVV  VLLPLVSSQ  LLPLVSSQC  LLALHRSYL  YLQPRTFLL  VLLPLVSSQ  LLPLVSSQC  TLDSKTQSL  LLALHRSYL | 517  512  502  6  7  269  151  109  119  6  7  241  269  6  7  109  241 |
| 4)COMPRED | HLA-A*02:06  HLA-A*03:01  HLA-A*11:01 | FVFLVLLPL  VLLPLVSSQ  LLPLVSSQC  TLDSKTQSL  IVNNATNVV  VLLPLVSSQ  LLPLVSSQC  LLALHRSYL  YLQPRTFLL  RLFRKSNLK  VLLPLVSSQ  LLPLVSSQC  TLDSKTQSL  LLALHRSYL  GVYFASTEK | 2  6  7  109  119  6  7  241  269  454  6  7  109  241  89 |
| IEDB T CELL EPITOPE PREDICTION TOOL | HLA-A*01:01  HLA-A*02:06  HLA-A*03:01  HLA-A*11:01  HLA-A*29:02  HLA-A*68:01  HLA-B*07:02  HLA-B*18:01  HLA-B*35:01  HLA-B*35:03  HLA-B*51:01  HLA-B*57:01  HLA-B*58:01  HLA-C*04:01  HLA-C*07:01  HLA-C*07:02 | LTDEMIAQY  WTAGAAAYY  TSNQVAVLY  CVADYSVLY  NIDGYFKIY  GAEHVNNSY  FQFCNDPFL  YQDVNCTEV  KQLSSNFGA  FVFLVLLPL  VVFLHVTYV  YLQPRTFLL  KLPDDFTGC  RLFRKSNLK  GVYYHKNNK  VTYVPAQEK  QIYKTPPIK  TLKSFTVEK  TLADAGFIK  QIAPGQTGK  EILPVSMTK  GVYFASTEK  SSTASALGK  RLFRKSNLK  VTYVPAQEK  NSASFSTFK  GVYYHKNNK  TLADAGFIK  GVLTESNKK  CVADYSVLY  WTAGAAAYY  WFVTQRNFY  VGGNYNYLY  YFPLQSYGF  NSASFSTFK  FASVYAWNR  FVIRGDEVR  NASVVNIQK  GTHWFVTQR  YNYLYRLFR  EILPVSMTK  SVYAWNRKR  APRDGQAYV  LPPAYTNSF  SPGSASSVA  IPTNFTISV  RAAEIRASA  MIAQYTSAL  TPINLVRDL  GPKKSTNLV  FPQSAPHGV  APGQTGKIA  IPIGAGICA  GEWVLLSTF  DEMIAQYTS  TESIVRFPN  MESEFRVYS  YEQGSGYIP  YENQKLIAN  NDLCFTNVY  IPFAMQMAY  SEFRVYSSA  FELLHAPAT  IPFAMQMAY  FAMQMAYRF  LPFNDGVYF  LPPAYTNSF  LPPLLTDEM  NATRFASVY  CVADYSVLY  FVSNGTHWF  NGVEGFNCY  FEYVSQPFL  FKNLREFVF  FPNITNLCP  FPLQSYGFQ  FVFLVLLPL  QPTESIVRF  FPREGVFVS  LPPAYTNSF  FTNVYADSF  DPPEAEVQI  LPLVSSQCV  IAIPTNFTI  LPFFSNVTW  FPQSAPHGV  LPDDFTGCV  LPPLLTDEM  DPPEAEVQI  LPFNDGVYF  LPPAYTNSF  PPLLTDEMI  IAPGQTGKI  RSFIEDLLF  LAGTITSGW  HADQLTPTW  HSAWSHPQF  TRFASVYAW  QSAPHGVVF  HADQLTPTW  IAIPTNFTI  RSFIEDLLF  LAGTITSGW  YSSANNCTF  HSAWSHPQF  GTITSGWTF  TSNQVAVLY  FAMQMAYRF  QSAPHGVVF  VFKNIDGYF  RFDNPVLPF  VYDPLQPEL  GFNCYFPLQ  VFQTRAGCL  QFNSAIGKI  YFPLQSYGF  TFGAGAALQ  RKDGEWVLL  RFQTLLALH  NSFTRGVYY  YRFNGIGVT  KRFDNPVLP  VRFPNITNL  IRGDEVRQI  VRDPQTLEI  TRFQTLLAL  FRSSVLHST  VVFLHVTYV  TRFASVYAW  VYDPLQPEL  FYEPQIITT  VRFPNITNL  NYNYLYRLF  EYVSQPFLM  YYPDKVFRS  FRKSNLKPF  VRKDGEWVL  QYGSFCTQL  VGYLQPRTF | 865  258  604  361  196  652  133  612  964  2  1060  269  424  454  142  1065  787  302  827  409  725  89  939  454  1065  370  142  827  550  361  258  1102  445  489  370  347  400  1173  1099  449  725  349  1216  24  680  714  1014  869  208  526  1052  411  664  1228  867  323  153  1206  917  388  896  155  515  896  898  84  24  861  343  361  1095  481  168  186  329  490  2  321  1089  24  392  985  8  712  56  1052  425  861  985  84  24  862  410  815  878  625  1257  345  1054  625  712  815  878  160  1257  880  604  898  1054  193  78  1137  485  642  926  489  887  1225  237  30  904  77  327  402  576  236  43  1060  345  1137  1109  327  448  169  37  456  1224  755  267 |
| 6)EPIJEN | HLA-A*01:01  HLA-A*02:06  HLA-A*03:01  HLA-A*11:01  HLA-A*68:01  HLA-B*35:01 | GAEHVNNSY  NIDGYFKIY  GTNTSNQVA  STECSNLLL  GIGVTQNVL  RISNCVADY  KTSVDCTMY  GIGVTQNVL  VIGIVNNTV  AISSVLNDI  RLQSLQTYV  VVFLHVTYV  FIEDLLFNK  RLFRKSNLK  TLADAGFIK  EILPVSMTK  QIYKTPPIK  QIAPGQTGK  VTYVPAQEK  GIYQTSNFR  FIEDLLFNK  GVLTESNKK  GVYFASTEK  GVYYHKNNK  VTYVPAQEK  EILPVSMTK  IADTTDAVR  FASVYAWNR  FGEVFNATR  FVIRGDEVR  RLFRKSNLK  QIAPGQTGK  FAMQMAYRF  LPFNDGVYF  LPPAYTNSF  TPINLVRDL  IPFAMQMAY  CPFGEVFNA  FVSNGTHWF  SVLYNSASF | 652  196  601  746  357  372  733  908  1129  972  1000  1060  817  454  827  725  787  409  1065  311  817  550  89  142  1065  725  569  347  338  400  454  409  898  84  24  208  896  336  1095  366 |
| 7)NETPANMHC 4.1 | HLA-A*01:01  HLA-A*03:01  HLA-B*07:02  HLA-B*58:01 | LTDEMIAQY  TSNQVAVLY  WTAGAAAYY  GAEHVNNSY  CVADYSVLY  KTSVDCTMY  NIDGYFKIY  ASFSTFKCY  RLFRKSNLK  QIYKTPPIK  VTYVPAQEK  TLKSFTVEK  QIAPGQTGK  APRDGQAYV  GPKKSTNLV  TPINLVRDL  FPQSAPHGV  QPTESIVRF  IPTNFTISV  LPPAYTNSF  SPGSASSVA  MIAQYTSAL  LPFNDGVYF  KPFERDIST  APGQTGKIA  RAAEIRASA  TLDSKTQSL  HADQLTPTW  RSFIEDLLF  LAGTITSGW  QSAPHGVVF  HSAWSHPQF  GTITSGWTF  TSNQVAVLY  IAIPTNFTI  FAMQMAYRF  YSSANNCTF | 865  604  258  652  361  733  196  372  454  787  1065  302  409  1216  526  208  1052  321  714  24  680  869  84  462  411  1014  109  625  815  878  1054  1257  880  604  712  898  160 |

**Table S2.** List showing the percentage of individuals within the epitope hits along with their cumulative percent.

**MHC-I Coverage in Indian Population:**

| **Number of epitope hits / HLA combinations recognized** | **Percent of individuals** | **Cumulative percent of population coverage** |
| --- | --- | --- |
| 0 | 16.92 | 100.0 |
| 1 | 0.0 | 83.08 |
| 2 | 0.0 | 83.08 |
| 3 | 0.0 | 83.08 |
| 4 | 0.0 | 83.08 |
| 5 | 0.0 | 83.08 |
| 6 | 0.0 | 83.08 |
| 7 | 0.0 | 83.08 |
| 8 | 0.0 | 83.08 |
| 9 | 0.0 | 83.08 |
| 10 | 16.31 | 83.08 |
| 11 | 2.53 | 66.77 |
| 12 | 0.0 | 64.23 |
| 13 | 0.0 | 64.23 |
| 14 | 4.63 | 64.23 |
| 15 | 7.59 | 59.6 |
| 16 | 0.0 | 52.01 |
| 17 | 0.0 | 52.01 |
| 18 | 0.0 | 52.01 |
| 19 | 3.99 | 52.01 |
| 20 | 5.01 | 48.02 |
| 21 | 2.01 | 43.01 |
| 22 | 0.04 | 41.0 |
| 23 | 1.64 | 40.96 |
| 24 | 4.19 | 39.33 |
| 25 | 7.81 | 35.13 |
| 26 | 1.14 | 27.32 |
| 27 | 0.0 | 26.18 |
| 28 | 0.26 | 26.18 |
| 29 | 4.97 | 25.92 |
| 30 | 1.49 | 20.95 |
| 31 | 0.43 | 19.46 |
| 32 | 0.02 | 19.03 |
| 33 | 2.27 | 19.01 |
| 34 | 2.36 | 16.74 |
| 35 | 2.6 | 14.37 |
| 36 | 0.9 | 11.77 |
| 37 | 0.29 | 10.88 |
| 38 | 0.66 | 10.58 |
| 39 | 2.18 | 9.92 |
| 40 | 0.92 | 7.74 |
| 41 | 0.09 | 6.82 |
| 42 | 0.21 | 6.73 |
| 43 | 1.12 | 6.52 |
| 44 | 1.27 | 5.4 |
| 45 | 0.46 | 4.13 |
| 46 | 0.19 | 3.67 |
| 47 | 0.28 | 3.49 |
| 48 | 0.58 | 3.21 |
| 49 | 0.48 | 2.63 |
| 50 | 0.28 | 2.15 |
| 51 | 0.06 | 1.87 |
| 52 | 0.23 | 1.81 |
| 53 | 0.29 | 1.58 |
| 54 | 0.35 | 1.29 |
| 55 | 0.11 | 0.94 |
| 56 | 0.03 | 0.83 |
| 57 | 0.1 | 0.8 |
| 58 | 0.21 | 0.7 |
| 59 | 0.09 | 0.49 |
| 60 | 0.03 | 0.4 |
| 61 | 0.02 | 0.37 |
| 62 | 0.09 | 0.34 |
| 63 | 0.07 | 0.26 |
| 64 | 0.04 | 0.19 |
| 65 | 0.02 | 0.15 |
| 66 | 0.02 | 0.13 |
| 67 | 0.02 | 0.11 |
| 68 | 0.03 | 0.09 |
| 69 | 0.01 | 0.06 |
| 70 | 0.0 | 0.05 |
| 71 | 0.01 | 0.05 |
| 72 | 0.01 | 0.04 |
| 73 | 0.01 | 0.03 |
| 74 | 0.0 | 0.02 |
| 75 | 0.0 | 0.01 |
| 76 | 0.0 | 0.01 |
| 77 | 0.0 | 0.01 |
| 78 | 0.0 | 0.01 |
| 79 | 0.0 | 0.0 |
| 80 | 0.0 | 0.0 |
| 81 | 0.0 | 0.0 |
| 82 | 0.0 | 0.0 |
| 83 | 0.0 | 0.0 |
| 84 | 0.0 | 0.0 |
| 85 | 0.0 | 0.0 |
| 86 | 0.0 | 0.0 |
| 87 | 0.0 | 0.0 |
| 88 | 0.0 | 0.0 |
| 89 | 0.0 | 0.0 |
| 90 | 0.0 | 0.0 |
| 91 | 0.0 | 0.0 |
| 92 | 0.0 | 0.0 |
| 93 | 0.0 | 0.0 |
| 94 | 0.0 | 0.0 |
| 95 | 0.0 | 0.0 |

**Table S3:** Coverage of individual epitopes in Indian population.

| **Epitope** | **Coverage** | **HLA allele (genotypic frequency (%))** | | | | | | | | | | | | | | | | | **Total HLA hits** |
| --- | --- | --- | --- | --- | --- | --- | --- | --- | --- | --- | --- | --- | --- | --- | --- | --- | --- | --- | --- |
|  | **Class I** | **HLA-A*01:01 (8.36)** | **HLA-A*02:06 (4.14)** | **HLA-A*03:01 (8.12)** | **HLA-A*11:01 (12.31)** | **HLA-A*29:02 (1.04)** | **HLA-A*68:01 (6.27)** | **HLA-B*07:02 (2.29)** | **HLA-B*18:01 (2.10)** | **HLA-B*35:01 (5.03)** | **HLA-B*35:03 (6.54)** | **HLA-B*50:01 (2.51)** | **HLA-B*51:01 (6.91)** | **HLA-B*57:01 (4.12)** | **HLA-B*58:01 (7.72)** | **HLA-C*04:01 (12.59)** | **HLA-C*07:01 (8.09)** | **HLA-C*07:02 (8.01)** |  |
| Epitope #1: ESEFRVYSS | 12.65% | + | - | - | - | - | - | - | - | - | - | - | - | - | - | - | - | - | **1** |
| Epitope #2: RLDPPEAEV | 12.65% | + | - | - | - | - | - | - | - | - | - | - | - | - | - | - | - | - | **1** |
| Epitope #3: NIDGYFKIY | 12.65% | + | - | - | - | - | - | - | - | - | - | - | - | - | - | - | - | - | **1** |
| Epitope #4: ECDIPIGAG | 12.65% | + | - | - | - | - | - | - | - | - | - | - | - | - | - | - | - | - | **1** |
| Epitope #5: LTDEMIAQY | 12.65% | + | - | - | - | - | - | - | - | - | - | - | - | - | - | - | - | - | **1** |
| Epitope #6: TSNQVAVLY | 12.65% | + | - | - | - | - | - | - | - | - | - | - | - | - | - | - | - | - | **1** |
| Epitope #7: WTAGAAAYY | 12.65% | + | - | - | - | - | - | - | - | - | - | - | - | - | - | - | - | - | **1** |
| Epitope #8: GAEHVNNSY | 12.65% | + | - | - | - | - | - | - | - | - | - | - | - | - | - | - | - | - | **1** |
| Epitope #9: CVADYSVLY | 12.65% | + | - | - | - | - | - | - | - | - | - | - | - | - | - | - | - | - | **1** |
| Epitope #10: RISNCVADY | 12.65% | + | - | - | - | - | - | - | - | - | - | - | - | - | - | - | - | - | **1** |
| Epitope #11: QGPGHHHHH | 12.65% | + | - | - | - | - | - | - | - | - | - | - | - | - | - | - | - | - | **1** |
| Epitope #12: KTSVDCTMY | 12.65% | + | - | - | - | - | - | - | - | - | - | - | - | - | - | - | - | - | **1** |
| Epitope #13: ASFSTFKCY | 12.65% | + | - | - | - | - | - | - | - | - | - | - | - | - | - | - | - | - | **1** |
| Epitope #14: ERDISTEIY | 12.65% | + | - | - | - | - | - | - | - | - | - | - | - | - | - | - | - | - | **1** |
| Epitope #15: NSFTRGVYY | 12.65% | + | - | - | - | - | - | - | - | - | - | - | - | - | - | - | - | - | **1** |
| Epitope #16: GTNTSNQVA | 12.65% | + | - | - | - | - | - | - | - | - | - | - | - | - | - | - | - | - | **1** |
| Epitope #17: STECSNLLL | 12.65% | + | - | - | - | - | - | - | - | - | - | - | - | - | - | - | - | - | **1** |
| Epitope #18: GIGVTQNVL | 12.65% | + | - | - | - | - | - | - | - | - | - | - | - | - | - | - | - | - | **1** |
| Epitope #19: IGAGICASY | 12.65% | + | - | - | - | - | - | - | - | - | - | - | - | - | - | - | - | - | **1** |
| Epitope #20: FQFCNDPFL | 6.37% | - | + | - | - | - | - | - | - | - | - | - | - | - | - | - | - | - | **1** |
| Epitope #21: YQDVNCTEV | 6.37% | - | + | - | - | - | - | - | - | - | - | - | - | - | - | - | - | - | **1** |
| Epitope #22: KQLSSNFGA | 6.37% | - | + | - | - | - | - | - | - | - | - | - | - | - | - | - | - | - | **1** |
| Epitope #23: FVFLVLLPL | 6.37% | - | + | - | - | - | - | - | - | - | - | - | - | - | - | - | - | - | **1** |
| Epitope #24: VVFLHVTYV | 6.37% | - | + | - | - | - | - | - | - | - | - | - | - | - | - | - | - | - | **1** |
| Epitope #25: YLQPRTFLL | 6.37% | - | + | - | - | - | - | - | - | - | - | - | - | - | - | - | - | - | **1** |
| Epitope #26: LQSYGFQPT | 6.37% | - | + | - | - | - | - | - | - | - | - | - | - | - | - | - | - | - | **1** |
| Epitope #27: KLPDDFTGC | 6.37% | - | + | - | - | - | - | - | - | - | - | - | - | - | - | - | - | - | **1** |
| Epitope #28: VIGIVNNTV | 6.37% | - | + | - | - | - | - | - | - | - | - | - | - | - | - | - | - | - | **1** |
| Epitope #29: AISSVLNDI | 6.37% | - | + | - | - | - | - | - | - | - | - | - | - | - | - | - | - | - | **1** |
| Epitope #30: RLQSLQTYV | 6.37% | - | + | - | - | - | - | - | - | - | - | - | - | - | - | - | - | - | **1** |
| Epitope #31: ELLHAPATV | 6.37% | - | + | - | - | - | - | - | - | - | - | - | - | - | - | - | - | - | **1** |
| Epitope #32: KIADYNYKL | 6.37% | - | + | - | - | - | - | - | - | - | - | - | - | - | - | - | - | - | **1** |
| Epitope #33: LLHAPATVC | 6.37% | - | + | - | - | - | - | - | - | - | - | - | - | - | - | - | - | - | **1** |
| Epitope #34: VLSFELLHA | 6.37% | - | + | - | - | - | - | - | - | - | - | - | - | - | - | - | - | - | **1** |
| Epitope #35: GVGYQPYRV | 6.37% | - | + | - | - | - | - | - | - | - | - | - | - | - | - | - | - | - | **1** |
| Epitope #36: VLLPLVSSQ | 6.37% | - | + | - | - | - | - | - | - | - | - | - | - | - | - | - | - | - | **1** |
| Epitope #37: LLPLVSSQC | 6.37% | - | + | - | - | - | - | - | - | - | - | - | - | - | - | - | - | - | **1** |
| Epitope #38: YLQPRTFLL | 6.37% | - | + | - | - | - | - | - | - | - | - | - | - | - | - | - | - | - | **1** |
| Epitope #39: SWMESEFRV | 6.37% | - | + | - | - | - | - | - | - | - | - | - | - | - | - | - | - | - | **1** |
| Epitope #40: TLDSKTQSL | 6.37% | - | + | - | - | - | - | - | - | - | - | - | - | - | - | - | - | - | **1** |
| Epitope #41: IVNNATNVV | 6.37% | - | + | - | - | - | - | - | - | - | - | - | - | - | - | - | - | - | **1** |
| Epitope #42: FVFLVLLPL | 6.37% | - | + | - | - | - | - | - | - | - | - | - | - | - | - | - | - | - | **1** |
| Epitope #43: FIEDLLFNK | 12.30% | - | - | + | - | - | - | - | - | - | - | - | - | - | - | - | - | - | **1** |
| Epitope #44: RLFRKSNLK | 12.30% | - | - | + | - | - | - | - | - | - | - | - | - | - | - | - | - | - | **1** |
| Epitope #45: TLADAGFIK | 12.30% | - | - | + | - | - | - | - | - | - | - | - | - | - | - | - | - | - | **1** |
| Epitope #46: EILPVSMTK | 12.30% | - | - | + | - | - | - | - | - | - | - | - | - | - | - | - | - | - | **1** |
| Epitope #47: QIYKTPPIK | 12.30% | - | - | + | - | - | - | - | - | - | - | - | - | - | - | - | - | - | **1** |
| Epitope #48: VTYVPAQEK | 12.30% | - | - | + | - | - | - | - | - | - | - | - | - | - | - | - | - | - | **1** |
| Epitope #49: GIYQTSNFR | 12.30% | - | - | + | - | - | - | - | - | - | - | - | - | - | - | - | - | - | **1** |
| Epitope #50: GVYYHKNNK | 12.30% | - | - | + | - | - | - | - | - | - | - | - | - | - | - | - | - | - | **1** |
| Epitope #51: TLKSFTVEK | 12.30% | - | - | + | - | - | - | - | - | - | - | - | - | - | - | - | - | - | **1** |
| Epitope #52: QIAPGQTGK | 12.30% | - | - | + | - | - | - | - | - | - | - | - | - | - | - | - | - | - | **1** |
| Epitope #53: VLLPLVSSQ | 12.30% | - | - | + | - | - | - | - | - | - | - | - | - | - | - | - | - | - | **1** |
| Epitope #54: LLPLVSSQC | 12.30% | - | - | + | - | - | - | - | - | - | - | - | - | - | - | - | - | - | **1** |
| Epitope #55: LLALHRSYL | 12.30% | - | - | + | - | - | - | - | - | - | - | - | - | - | - | - | - | - | **1** |
| Epitope #56: YLQPRTFLL | 12.30% | - | - | + | - | - | - | - | - | - | - | - | - | - | - | - | - | - | **1** |
| Epitope #57: APRDGQAYV | 2.74% | - | - | - | - | - | - | + | - | - | - | - | - | - | - | - | - | - | **1** |
| Epitope #58: GPKKSTNLV | 2.74% | - | - | - | - | - | - | + | - | - | - | - | - | - | - | - | - | - | **1** |
| Epitope #59: TPINLVRDL | 2.74% | - | - | - | - | - | - | + | - | - | - | - | - | - | - | - | - | - | **1** |
| Epitope #60: FPQSAPHGV | 2.74% | - | - | - | - | - | - | + | - | - | - | - | - | - | - | - | - | - | **1** |
| Epitope #61: QPTESIVRF | 2.74% | - | - | - | - | - | - | + | - | - | - | - | - | - | - | - | - | - | **1** |
| Epitope #62: IPTNFTISV | 2.74% | - | - | - | - | - | - | + | - | - | - | - | - | - | - | - | - | - | **1** |
| Epitope #63: LPPAYTNSF | 2.74% | - | - | - | - | - | - | + | - | - | - | - | - | - | - | - | - | - | **1** |
| Epitope #64: SPGSASSVA | 2.74% | - | - | - | - | - | - | + | - | - | - | - | - | - | - | - | - | - | **1** |
| Epitope #65: MIAQYTSAL | 2.74% | - | - | - | - | - | - | + | - | - | - | - | - | - | - | - | - | - | **1** |
| Epitope #66: LPFNDGVYF | 2.74% | - | - | - | - | - | - | + | - | - | - | - | - | - | - | - | - | - | **1** |
| Epitope #67: KPFERDIST | 2.74% | - | - | - | - | - | - | + | - | - | - | - | - | - | - | - | - | - | **1** |
| Epitope #68: APGQTGKIA | 2.74% | - | - | - | - | - | - | + | - | - | - | - | - | - | - | - | - | - | **1** |
| Epitope #69: RAAEIRASA | 2.74% | - | - | - | - | - | - | + | - | - | - | - | - | - | - | - | - | - | **1** |
| Epitope #70: TLDSKTQSL | 2.74% | - | - | - | - | - | - | + | - | - | - | - | - | - | - | - | - | - | **1** |
| Epitope #71: IPFAMQMAY | 2.74% | - | - | - | - | - | - | + | - | - | - | - | - | - | - | - | - | - | **1** |
| Epitope #72: FPQSAPHGV | 2.74% | - | - | - | - | - | - | + | - | - | - | - | - | - | - | - | - | - | **1** |
| Epitope #73: IPIGAGICA | 2.74% | - | - | - | - | - | - | + | - | - | - | - | - | - | - | - | - | - | **1** |
| Epitope #74: CPFGEVFNA | 2.74% | - | - | - | - | - | - | + | - | - | - | - | - | - | - | - | - | - | **1** |
| Epitope #75: LPDDFTGCV | 2.74% | - | - | - | - | - | - | + | - | - | - | - | - | - | - | - | - | - | **1** |
| Epitope #76: LPPLLTDEM | 5.98% | - | - | - | - | - | - | - | - | + | - | - | - | - | - | - | - | - | **1** |
| Epitope #77: LPFFSNVTW | 5.98% | - | - | - | - | - | - | - | - | + | - | - | - | - | - | - | - | - | **1** |
| Epitope #78: IPFAMQMAY | 5.98% | - | - | - | - | - | - | - | - | + | - | - | - | - | - | - | - | - | **1** |
| Epitope #79: TPINLVRDL | 5.98% | - | - | - | - | - | - | - | - | + | - | - | - | - | - | - | - | - | **1** |
| Epitope #80: TPCNGVEGF | 5.98% | - | - | - | - | - | - | - | - | + | - | - | - | - | - | - | - | - | **1** |
| Epitope #81: LPPAYTNSF | 5.98% | - | - | - | - | - | - | - | - | + | - | - | - | - | - | - | - | - | **1** |
| Epitope #82: FAMQMAYRF | 5.98% | - | - | - | - | - | - | - | - | + | - | - | - | - | - | - | - | - | **1** |
| Epitope #83: LPFNDGVYF | 5.98% | - | - | - | - | - | - | - | - | + | - | - | - | - | - | - | - | - | **1** |
| Epitope #84: FVSNGTHWF | 5.98% | - | - | - | - | - | - | - | - | + | - | - | - | - | - | - | - | - | **1** |
| Epitope #85: CPFGEVFNA | 5.98% | - | - | - | - | - | - | - | - | + | - | - | - | - | - | - | - | - | **1** |
| Epitope #86: SVLYNSASF | 5.98% | - | - | - | - | - | - | - | - | + | - | - | - | - | - | - | - | - | **1** |
| Epitope #87: NATRFASVY | 5.98% | - | - | - | - | - | - | - | - | + | - | - | - | - | - | - | - | - | **1** |
| Epitope #88: NGVEGFNCY | 5.98% | - | - | - | - | - | - | - | - | + | - | - | - | - | - | - | - | - | **1** |
| Epitope #89: CVADYSVLY | 5.98% | - | - | - | - | - | - | - | - | + | - | - | - | - | - | - | - | - | **1** |
| Epitope #90: DPPEAEVQI | 8.16% | - | - | - | - | - | - | - | - | - | - | - | + | - | - | - | - | - | **1** |
| Epitope #91: IAIPTNFTI | 8.16% | - | - | - | - | - | - | - | - | - | - | - | + | - | - | - | - | - | **1** |
| Epitope #92: LPPLLTDEM | 8.16% | - | - | - | - | - | - | - | - | - | - | - | + | - | - | - | - | - | **1** |
| Epitope #93: LPPAYTNSF | 8.16% | - | - | - | - | - | - | - | - | - | - | - | + | - | - | - | - | - | **1** |
| Epitope #94: LPLVSSQCV | 8.16% | - | - | - | - | - | - | - | - | - | - | - | + | - | - | - | - | - | **1** |
| Epitope #95: PPLLTDEMI | 8.16% | - | - | - | - | - | - | - | - | - | - | - | + | - | - | - | - | - | **1** |
| Epitope #96: IAPGQTGKI | 8.16% | - | - | - | - | - | - | - | - | - | - | - | + | - | - | - | - | - | **1** |
| Epitope #97: FPQSAPHGV | 8.16% | - | - | - | - | - | - | - | - | - | - | - | + | - | - | - | - | - | **1** |
| Epitope #98: LPFNDGVYF | 8.16% | - | - | - | - | - | - | - | - | - | - | - | + | - | - | - | - | - | **1** |
| Epitope #99: LPFFSNVTW | 8.16% | - | - | - | - | - | - | - | - | - | - | - | + | - | - | - | - | - | **1** |
| Epitope #100: LPDDFTGCV | 8.16% | - | - | - | - | - | - | - | - | - | - | - | + | - | - | - | - | - | **1** |
| Epitope #101: GEVFNATRF | 1.62% | - | - | - | - | + | - | - | - | - | - | - | - | - | - | - | - | - | **1** |
| Epitope #102: AEIRASANL | 1.62% | - | - | - | - | + | - | - | - | - | - | - | - | - | - | - | - | - | **1** |
| Epitope #103: QEVFAQVKQ | 1.62% | - | - | - | - | + | - | - | - | - | - | - | - | - | - | - | - | - | **1** |
| Epitope #104: FERDISTEI | 1.62% | - | - | - | - | + | - | - | - | - | - | - | - | - | - | - | - | - | **1** |
| Epitope #105: SKVGGNYNY | 1.62% | - | - | - | - | + | - | - | - | - | - | - | - | - | - | - | - | - | **1** |
| Epitope #106: CVADYSVLY | 1.62% | - | - | - | - | + | - | - | - | - | - | - | - | - | - | - | - | - | **1** |
| Epitope #107: YFPLQSYGF | 1.62% | - | - | - | - | + | - | - | - | - | - | - | - | - | - | - | - | - | **1** |
| Epitope #108: VGGNYNYLY | 1.62% | - | - | - | - | + | - | - | - | - | - | - | - | - | - | - | - | - | **1** |
| Epitope #109: WTAGAAAYY | 1.62% | - | - | - | - | + | - | - | - | - | - | - | - | - | - | - | - | - | **1** |
| Epitope #110: WFVTQRNFY | 1.62% | - | - | - | - | + | - | - | - | - | - | - | - | - | - | - | - | - | **1** |
| Epitope #111: GVYFASTEK | 18.32% | - | - | - | + | - | - | - | - | - | - | - | - | - | - | - | - | - | **1** |
| Epitope #112: GVYYHKNNK | 18.32% | - | - | - | + | - | - | - | - | - | - | - | - | - | - | - | - | - | **1** |
| Epitope #113: RLFRKSNLK | 18.32% | - | - | - | + | - | - | - | - | - | - | - | - | - | - | - | - | - | **1** |
| Epitope #114: EILPVSMTK | 18.32% | - | - | - | + | - | - | - | - | - | - | - | - | - | - | - | - | - | **1** |
| Epitope #115: TLADAGFIK | 18.32% | - | - | - | + | - | - | - | - | - | - | - | - | - | - | - | - | - | **1** |
| Epitope #116: FIEDLLFNK | 18.32% | - | - | - | + | - | - | - | - | - | - | - | - | - | - | - | - | - | **1** |
| Epitope #117: GVLTESNKK | 18.32% | - | - | - | + | - | - | - | - | - | - | - | - | - | - | - | - | - | **1** |
| Epitope #118: ASFSTFKCY | 18.32% | - | - | - | + | - | - | - | - | - | - | - | - | - | - | - | - | - | **1** |
| Epitope #119: VTYVPAQEK | 18.32% | - | - | - | + | - | - | - | - | - | - | - | - | - | - | - | - | - | **1** |
| Epitope #120: SSTASALGK | 18.32% | - | - | - | + | - | - | - | - | - | - | - | - | - | - | - | - | - | **1** |
| Epitope #121: NSASFSTFK | 18.32% | - | - | - | + | - | - | - | - | - | - | - | - | - | - | - | - | - | **1** |
| Epitope #122: VLLPLVSSQ | 18.32% | - | - | - | + | - | - | - | - | - | - | - | - | - | - | - | - | - | **1** |
| Epitope #123: LLPLVSSQC | 18.32% | - | - | - | + | - | - | - | - | - | - | - | - | - | - | - | - | - | **1** |
| Epitope #124: TLDSKTQSL | 18.32% | - | - | - | + | - | - | - | - | - | - | - | - | - | - | - | - | - | **1** |
| Epitope #125: LLALHRSYL | 18.32% | - | - | - | + | - | - | - | - | - | - | - | - | - | - | - | - | - | **1** |
| Epitope #126: GTHWFVTQR | 9.56% | - | - | - | - | - | + | - | - | - | - | - | - | - | - | - | - | - | **1** |
| Epitope #127: SVYAWNRKR | 9.56% | - | - | - | - | - | + | - | - | - | - | - | - | - | - | - | - | - | **1** |
| Epitope #128: QIAPGQTGK | 9.56% | - | - | - | - | - | + | - | - | - | - | - | - | - | - | - | - | - | **1** |
| Epitope #129: IADTTDAVR | 9.56% | - | - | - | - | - | + | - | - | - | - | - | - | - | - | - | - | - | **1** |
| Epitope #130: NASVVNIQK | 9.56% | - | - | - | - | - | + | - | - | - | - | - | - | - | - | - | - | - | **1** |
| Epitope #131: FVIRGDEVR | 9.56% | - | - | - | - | - | + | - | - | - | - | - | - | - | - | - | - | - | **1** |
| Epitope #132: NSASFSTFK | 9.56% | - | - | - | - | - | + | - | - | - | - | - | - | - | - | - | - | - | **1** |
| Epitope #133: GVLTESNKK | 9.56% | - | - | - | - | - | + | - | - | - | - | - | - | - | - | - | - | - | **1** |
| Epitope #134: GVYFASTEK | 9.56% | - | - | - | - | - | + | - | - | - | - | - | - | - | - | - | - | - | **1** |
| Epitope #135: EILPVSMTK | 9.56% | - | - | - | - | - | + | - | - | - | - | - | - | - | - | - | - | - | **1** |
| Epitope #136: RLFRKSNLK | 9.56% | - | - | - | - | - | + | - | - | - | - | - | - | - | - | - | - | - | **1** |
| Epitope #137: SVYAWNRKR | 9.56% | - | - | - | - | - | + | - | - | - | - | - | - | - | - | - | - | - | **1** |
| Epitope #138: FASVYAWNR | 9.56% | - | - | - | - | - | + | - | - | - | - | - | - | - | - | - | - | - | **1** |
| Epitope #139: FGEVFNATR | 9.56% | - | - | - | - | - | + | - | - | - | - | - | - | - | - | - | - | - | **1** |
| Epitope #140: YNYLYRLFR | 9.56% | - | - | - | - | - | + | - | - | - | - | - | - | - | - | - | - | - | **1** |
| Epitope #141: KTPPIKDFG | 4.91% | - | - | - | - | - | - | - | - | - | - | - | - | + | - | - | - | - | **1** |
| Epitope #142: FTNVYADSF | 4.91% | - | - | - | - | - | - | - | - | - | - | - | - | + | - | - | - | - | **1** |
| Epitope #143: QSAPHGVVF | 4.91% | - | - | - | - | - | - | - | - | - | - | - | - | + | - | - | - | - | **1** |
| Epitope #144: HADQLTPTW | 4.91% | - | - | - | - | - | - | - | - | - | - | - | - | + | - | - | - | - | **1** |
| Epitope #145: DFTGCVIAW | 4.91% | - | - | - | - | - | - | - | - | - | - | - | - | + | - | - | - | - | **1** |
| Epitope #146: HHHHHHSAW | 4.91% | - | - | - | - | - | - | - | - | - | - | - | - | + | - | - | - | - | **1** |
| Epitope #147: RSFIEDLLF | 4.91% | - | - | - | - | - | - | - | - | - | - | - | - | + | - | - | - | - | **1** |
| Epitope #148: LAGTITSGW | 4.91% | - | - | - | - | - | - | - | - | - | - | - | - | + | - | - | - | - | **1** |
| Epitope #149: HSAWSHPQF | 4.91% | - | - | - | - | - | - | - | - | - | - | - | - | + | - | - | - | - | **1** |
| Epitope #150: TRFASVYAW | 4.91% | - | - | - | - | - | - | - | - | - | - | - | - | + | - | - | - | - | **1** |
| Epitope #151: HADQLTPTW | 9.10% | - | - | - | - | - | - | - | - | - | - | - | - | - | + | - | - | - | **1** |
| Epitope #152: IAIPTNFTI | 9.10% | - | - | - | - | - | - | - | - | - | - | - | - | - | + | - | - | - | **1** |
| Epitope #153: RSFIEDLLF | 9.10% | - | - | - | - | - | - | - | - | - | - | - | - | - | + | - | - | - | **1** |
| Epitope #154: LAGTITSGW | 9.10% | - | - | - | - | - | - | - | - | - | - | - | - | - | + | - | - | - | **1** |
| Epitope #155: YSSANNCTF | 9.10% | - | - | - | - | - | - | - | - | - | - | - | - | - | + | - | - | - | **1** |
| Epitope #156: HSAWSHPQF | 9.10% | - | - | - | - | - | - | - | - | - | - | - | - | - | + | - | - | - | **1** |
| Epitope #157: GTITSGWTF | 9.10% | - | - | - | - | - | - | - | - | - | - | - | - | - | + | - | - | - | **1** |
| Epitope #158: TSNQVAVLY | 9.10% | - | - | - | - | - | - | - | - | - | - | - | - | - | + | - | - | - | **1** |
| Epitope #159: FAMQMAYRF | 9.10% | - | - | - | - | - | - | - | - | - | - | - | - | - | + | - | - | - | **1** |
| Epitope #160: QSAPHGVVF | 9.10% | - | - | - | - | - | - | - | - | - | - | - | - | - | + | - | - | - | **1** |
| Epitope #161: SEFRVYSSA | 3.01% | - | - | - | - | - | - | - | - | - | - | + | - | - | - | - | - | - | **1** |
| Epitope #162: TEVPVAIHA | 3.01% | - | - | - | - | - | - | - | - | - | - | + | - | - | - | - | - | - | **1** |
| Epitope #163: NENGTITDA | 3.01% | - | - | - | - | - | - | - | - | - | - | + | - | - | - | - | - | - | **1** |
| Epitope #164: YECDIPIGA | 3.01% | - | - | - | - | - | - | - | - | - | - | + | - | - | - | - | - | - | **1** |
| Epitope #165: PEAPRDGQA | 3.01% | - | - | - | - | - | - | - | - | - | - | + | - | - | - | - | - | - | **1** |
| Epitope #166: FELLHAPAT | 3.01% | - | - | - | - | - | - | - | - | - | - | + | - | - | - | - | - | - | **1** |
| Epitope #167: RAAEIRASA | 3.01% | - | - | - | - | - | - | - | - | - | - | + | - | - | - | - | - | - | **1** |
| Epitope #168: TTRTQLPPA | 3.01% | - | - | - | - | - | - | - | - | - | - | + | - | - | - | - | - | - | **1** |
| Epitope #169: MESEFRVYS | 3.01% | - | - | - | - | - | - | - | - | - | - | + | - | - | - | - | - | - | **1** |
| Epitope #170: FEYVSQPFL | 3.01% | - | - | - | - | - | - | - | - | - | - | + | - | - | - | - | - | - | **1** |
| Epitope #171: RDLPQGFSA | 3.01% | - | - | - | - | - | - | - | - | - | - | + | - | - | - | - | - | - | **1** |
| Epitope #172: GEWVLLSTF | 2.52% | - | - | - | - | - | - | - | + | - | - | - | - | - | - | - | - | - | **1** |
| Epitope #173: DEMIAQYTS | 2.52% | - | - | - | - | - | - | - | + | - | - | - | - | - | - | - | - | - | **1** |
| Epitope #174: TESIVRFPN | 2.52% | - | - | - | - | - | - | - | + | - | - | - | - | - | - | - | - | - | **1** |
| Epitope #175: MESEFRVYS | 2.52% | - | - | - | - | - | - | - | + | - | - | - | - | - | - | - | - | - | **1** |
| Epitope #176: YEQGSGYIP | 2.52% | - | - | - | - | - | - | - | + | - | - | - | - | - | - | - | - | - | **1** |
| Epitope #177: YENQKLIAN | 2.52% | - | - | - | - | - | - | - | + | - | - | - | - | - | - | - | - | - | **1** |
| Epitope #178: NDLCFTNVY | 2.52% | - | - | - | - | - | - | - | + | - | - | - | - | - | - | - | - | - | **1** |
| Epitope #179: IPFAMQMAY | 2.52% | - | - | - | - | - | - | - | + | - | - | - | - | - | - | - | - | - | **1** |
| Epitope #180: SEFRVYSSA | 2.52% | - | - | - | - | - | - | - | + | - | - | - | - | - | - | - | - | - | **1** |
| Epitope #181: FELLHAPAT | 2.52% | - | - | - | - | - | - | - | + | - | - | - | - | - | - | - | - | - | **1** |
| Epitope #182: FEYVSQPFL | 7.73% | - | - | - | - | - | - | - | - | - | + | - | - | - | - | - | - | - | **1** |
| Epitope #183: FKNLREFVF | 7.73% | - | - | - | - | - | - | - | - | - | + | - | - | - | - | - | - | - | **1** |
| Epitope #184: FPNITNLCP | 7.73% | - | - | - | - | - | - | - | - | - | + | - | - | - | - | - | - | - | **1** |
| Epitope #185: FPLQSYGFQ | 7.73% | - | - | - | - | - | - | - | - | - | + | - | - | - | - | - | - | - | **1** |
| Epitope #186: FVFLVLLPL | 7.73% | - | - | - | - | - | - | - | - | - | + | - | - | - | - | - | - | - | **1** |
| Epitope #187: QPTESIVRF | 7.73% | - | - | - | - | - | - | - | - | - | + | - | - | - | - | - | - | - | **1** |
| Epitope #188: FPREGVFVS | 7.73% | - | - | - | - | - | - | - | - | - | + | - | - | - | - | - | - | - | **1** |
| Epitope #189: LPPAYTNSF | 7.73% | - | - | - | - | - | - | - | - | - | + | - | - | - | - | - | - | - | **1** |
| Epitope #190: FTNVYADSF | 7.73% | - | - | - | - | - | - | - | - | - | + | - | - | - | - | - | - | - | **1** |
| Epitope #191: DPPEAEVQI | 7.73% | - | - | - | - | - | - | - | - | - | + | - | - | - | - | - | - | - | **1** |
| Epitope #192: VFKNIDGYF | 18.85% | - | - | - | - | - | - | - | - | - | - | - | - | - | - | + | - | - | **1** |
| Epitope #193: RFDNPVLPF | 18.85% | - | - | - | - | - | - | - | - | - | - | - | - | - | - | + | - | - | **1** |
| Epitope #194: VYDPLQPEL | 18.85% | - | - | - | - | - | - | - | - | - | - | - | - | - | - | + | - | - | **1** |
| Epitope #195: GFNCYFPLQ | 18.85% | - | - | - | - | - | - | - | - | - | - | - | - | - | - | + | - | - | **1** |
| Epitope #196: VFQTRAGCL | 18.85% | - | - | - | - | - | - | - | - | - | - | - | - | - | - | + | - | - | **1** |
| Epitope #197: QFNSAIGKI | 18.85% | - | - | - | - | - | - | - | - | - | - | - | - | - | - | + | - | - | **1** |
| Epitope #198: YFPLQSYGF | 18.85% | - | - | - | - | - | - | - | - | - | - | - | - | - | - | + | - | - | **1** |
| Epitope #199: TFGAGAALQ | 18.85% | - | - | - | - | - | - | - | - | - | - | - | - | - | - | + | - | - | **1** |
| Epitope #200: RKDGEWVLL | 18.85% | - | - | - | - | - | - | - | - | - | - | - | - | - | - | + | - | - | **1** |
| Epitope #201: RFQTLLALH | 18.85% | - | - | - | - | - | - | - | - | - | - | - | - | - | - | + | - | - | **1** |
| Epitope #202: NSFTRGVYY | 12.34% | - | - | - | - | - | - | - | - | - | - | - | - | - | - | - | + | - | **1** |
| Epitope #203: YRFNGIGVT | 12.34% | - | - | - | - | - | - | - | - | - | - | - | - | - | - | - | + | - | **1** |
| Epitope #204: KRFDNPVLP | 12.34% | - | - | - | - | - | - | - | - | - | - | - | - | - | - | - | + | - | **1** |
| Epitope #205: VRFPNITNL | 12.34% | - | - | - | - | - | - | - | - | - | - | - | - | - | - | - | + | - | **1** |
| Epitope #206: IRGDEVRQI | 12.34% | - | - | - | - | - | - | - | - | - | - | - | - | - | - | - | + | - | **1** |
| Epitope #207: VRDPQTLEI | 12.34% | - | - | - | - | - | - | - | - | - | - | - | - | - | - | - | + | - | **1** |
| Epitope #208: TRFQTLLAL | 12.34% | - | - | - | - | - | - | - | - | - | - | - | - | - | - | - | + | - | **1** |
| Epitope #209: FRSSVLHST | 12.34% | - | - | - | - | - | - | - | - | - | - | - | - | - | - | - | + | - | **1** |
| Epitope #210: VVFLHVTYV | 12.34% | - | - | - | - | - | - | - | - | - | - | - | - | - | - | - | + | - | **1** |
| Epitope #211: TRFASVYAW | 12.34% | - | - | - | - | - | - | - | - | - | - | - | - | - | - | - | + | - | **1** |
| Epitope #212: VYDPLQPEL | 12.22% | - | - | - | - | - | - | - | - | - | - | - | - | - | - | - | - | + | **1** |
| Epitope #213: FYEPQIITT | 12.22% | - | - | - | - | - | - | - | - | - | - | - | - | - | - | - | - | + | **1** |
| Epitope #214: VRFPNITNL | 12.22% | - | - | - | - | - | - | - | - | - | - | - | - | - | - | - | - | + | **1** |
| Epitope #215: NYNYLYRLF | 12.22% | - | - | - | - | - | - | - | - | - | - | - | - | - | - | - | - | + | **1** |
| Epitope #216: EYVSQPFLM | 12.22% | - | - | - | - | - | - | - | - | - | - | - | - | - | - | - | - | + | **1** |
| Epitope #217: YYPDKVFRS | 12.22% | - | - | - | - | - | - | - | - | - | - | - | - | - | - | - | - | + | **1** |
| Epitope #218: FRKSNLKPF | 12.22% | - | - | - | - | - | - | - | - | - | - | - | - | - | - | - | - | + | **1** |
| Epitope #219: VRKDGEWVL | 12.22% | - | - | - | - | - | - | - | - | - | - | - | - | - | - | - | - | + | **1** |
| Epitope #220: QYGSFCTQL | 12.22% | - | - | - | - | - | - | - | - | - | - | - | - | - | - | - | - | + | **1** |
| Epitope #221: VGYLQPRTF | 12.22% | - | - | - | - | - | - | - | - | - | - | - | - | - | - | - | - | + | **1** |
| **Epitope set** | **83.08%** | **19** | **23** | **14** | **15** | **10** | **15** | **19** | **10** | **14** | **10** | **11** | **11** | **10** | **10** | **10** | **10** | **10** | **221** |

+ : restricted
- : not restricted
shaded column : genotypic frequency of this allele is 0 (zero)
